# Precision Confidence Mapping: An approach to determining individualized network topography with limited data

**DOI:** 10.64898/2026.08.17.744952

**Authors:** Julian S.B. Ramirez, Robert J.M. Hermosillo, Julia Moser, Gracie J. Grimsrud, Erendiz Tarakci, Han H. N. Pham, Kate J. Godfrey, Helayna Sjoberg, Vanessa Morgan, Thomas J. Madison, Timothy O. Laumann, Evan M. Gordon, Nico U.F. Dosenbach, Kimberly B. Weldon, Oscar Miranda-Dominguez, Brenden Tervo-Clemmens, Steven M. Nelson, Damien A. Fair

## Abstract

Individualized resting-state functional magnetic resonance imaging (rs-fMRI) is increasingly used to guide neuromodulation target selection. However, clinical scans are often short and noisy, and standard pipelines for functional network identification do not provide information about confidence of network assignment. With limited data, unstable network assignments can misdirect stimulation toward off-target regions, making it critical to know which assignments can be trusted. We developed Precision Confidence Mapping (PCM), a bootstrap-based framework that makes this uncertainty explicit and actionable. PCM repeatedly resamples the time series and reruns network detection to estimate how consistently each vertex is assigned to a given network. The resulting confidence maps can be thresholded to exclude less stable regions. We evaluated PCM across scan durations from 5 to 70 minutes using positive predictive value (PPV) as the primary measure of network-assignment precision. PPV quantified the proportion of vertices assigned to a network that received the same label in an independent within-subject 70 minute reference map. Confidence thresholding markedly improved PPV across functional networks, with the largest gains for short scan durations. Compared with standard network assignment, PCM significantly increased agreement with this independent reference. Within-subject agreement remained greater than between-subject agreement, indicating that thresholding preserved individual-specific network topography. These precision gains came with modest reductions in reference-network coverage, particularly at shorter scan durations. This tradeoff may be acceptable for neuromodulation applications that prioritize minimizing off-network assignments. By adding a reliability layer to individualized mapping, PCM supports more cautious and precise neuromodulation targeting under real-world clinical scan constraints.

## Introduction

Neuromodulation of the human brain has a long history in both therapeutic intervention and scientific discovery ^1–5^. Its use is now accelerating rapidly, driven by growing interest in alternatives to pharmacological therapies and by the emergence of modern noninvasive and invasive neuromodulation technologies ^6–13^.

A central limitation of neuromodulation approaches, particularly for psychiatric and other applications that target distributed association networks, is identifying the optimal stimulation target for a given individual. Historically, neuromodulation targets have often been defined using anatomical landmarks or group-average coordinates without accounting for individual functional network organization ^9,14,15^. Emerging evidence suggests that effective stimulation sites may be better understood in relation to functional networks and connectivity profiles than as isolated anatomical coordinates ^9,16–23^. However, the human cortex is highly variable in its functional organization across individuals; as such, target selection based on group-average localization may engage different circuits or miss the intended functional circuit entirely ^24–27^. This issue is especially pronounced in association cortex, where distributed functional networks vary substantially from person to person, such that otherwise similar anatomical locations in two individuals can belong to distinct functional systems ^26,28–31^. This variability may contribute to inconsistent efficacy in neuromodulation applications that target association cortex.

To address this targeting problem, investigators have increasingly turned to Precision Functional Mapping (PFM), which aims to identify individual-specific functional network topography rather than relying on group-average organization or imprecise anatomic descriptors. In resting-state applications, PFM leverages dense rs-fMRI sampling and community-detection methods to identify reliable person-specific network topography, thereby enabling subject-specific target selection ^24,25,32,33^. Individualized rs-fMRI-guided neuromodulation approaches are being used in both research and clinical settings and have shown encouraging early results across multiple applications ^3,19,34–38^. Individualized localization provides a principled path toward personalized neuromodulation targeting beyond one-size-fits-all anatomical rules. However, translating this approach to individual clinical use requires reliable network estimates given the quantity and quality of data acquired for each person.

In current studies using rs-fMRI to inform neuromodulation in individual patients, acquisitions are often brief, commonly 5 to 15 minutes. Resting-state fMRI has a low signal-to-noise ratio, and head motion and physiological noise reduce reliability ^39–42^. Individualized network mapping is feasible with dense-sampling approaches such as PFM. However, stable individual-level community detection may require substantially more data, in some cases exceeding 100 minutes^24^. Longer acquisitions may therefore be warranted when justified by the intended clinical application. With brief acquisitions, the spatial location of connectivity-defined targets, including those in dorsolateral prefrontal cortex, can vary across sessions, limiting their reproducibility^14^. Regardless of the amount of data collected, the uncertainty associated with the resulting network estimate should be characterized.

Beyond the effects of data quantity and quality, most community-detection approaches used to derive functional networks from resting-state fMRI do not quantify confidence in individual network assignments. Instead, they assign every vertex or voxel a single network label ^25,43^.

This hard assignment obscures both methodological and biological sources of ambiguity. First, network assignments can depend on the detection procedure and parameter choices, and stochastic algorithms such as Infomap can yield different solutions across runs ^44^. Second, brain regions do not always have a singular network identity ^30,45,46^. Some regions show meaningful affinity to multiple networks (e.g., see ^30^). Current targeting approaches rarely provide an explicit estimate of how confidently a given cortical or subcortical location belongs to the target network. Consequently, methods are needed to distinguish high-confidence network assignments from uncertain ones, so that targeting decisions can explicitly account for uncertainty, regardless of data quantity.

Here, we introduce Precision Confidence Mapping (PCM), a framework designed to make uncertainty in individualized network mapping explicit and actionable for neuromodulation targeting. Rather than producing a single, hard network assignment from a given rs-fMRI dataset, PCM repeatedly resamples the time series to generate multiple pseudo-replicates and applies a community-detection algorithm (e.g., template matching or Infomap) to each. Aggregating these solutions yields vertex-wise estimates of network assignment stability and probability, revealing where network membership is robust and where it is ambiguous. By providing individualized confidence maps instead of deterministic labels, PCM enables targeting strategies that explicitly account for uncertainty, prioritizing regions that are most likely to lie within the intended network when data are limited.

## Results

### Limited data reduce the precision of individualized networks

We first evaluated the accuracy of a standard community-detection method (i.e., template matching) applied to PFM-length (70 minute) versus traditional (5 minute) resting-state scans in four densely sampled participants. Figure 1A–B illustrates the tradeoff motivating this analysis. Group-average maps provide canonical network definitions but do not capture individual-specific network topography, while individualized mapping preserves this variation but requires sufficient data to produce precise assignments. For each participant, the resting-state data were divided into independent exploratory and reference halves (Fig. 1E). To define a within-subject ground truth reference, we derived network labels from an independent held-out 70 minute PFM segment in the reference half (Fig. 1B,E). We refer to this reference as the ground truth for brevity throughout the manuscript, although it should be interpreted as an independent benchmark rather than a fully converged estimate, because even 70 minutes of data may not completely stabilize vertex-wise functional connectivity or network assignment. Network assignments were then generated using template matching from either 5 or 70 minutes of data from the exploratory half and compared with the held-out ground truth to assess assignment accuracy (Fig. 1C,D).

**Figure 1:**
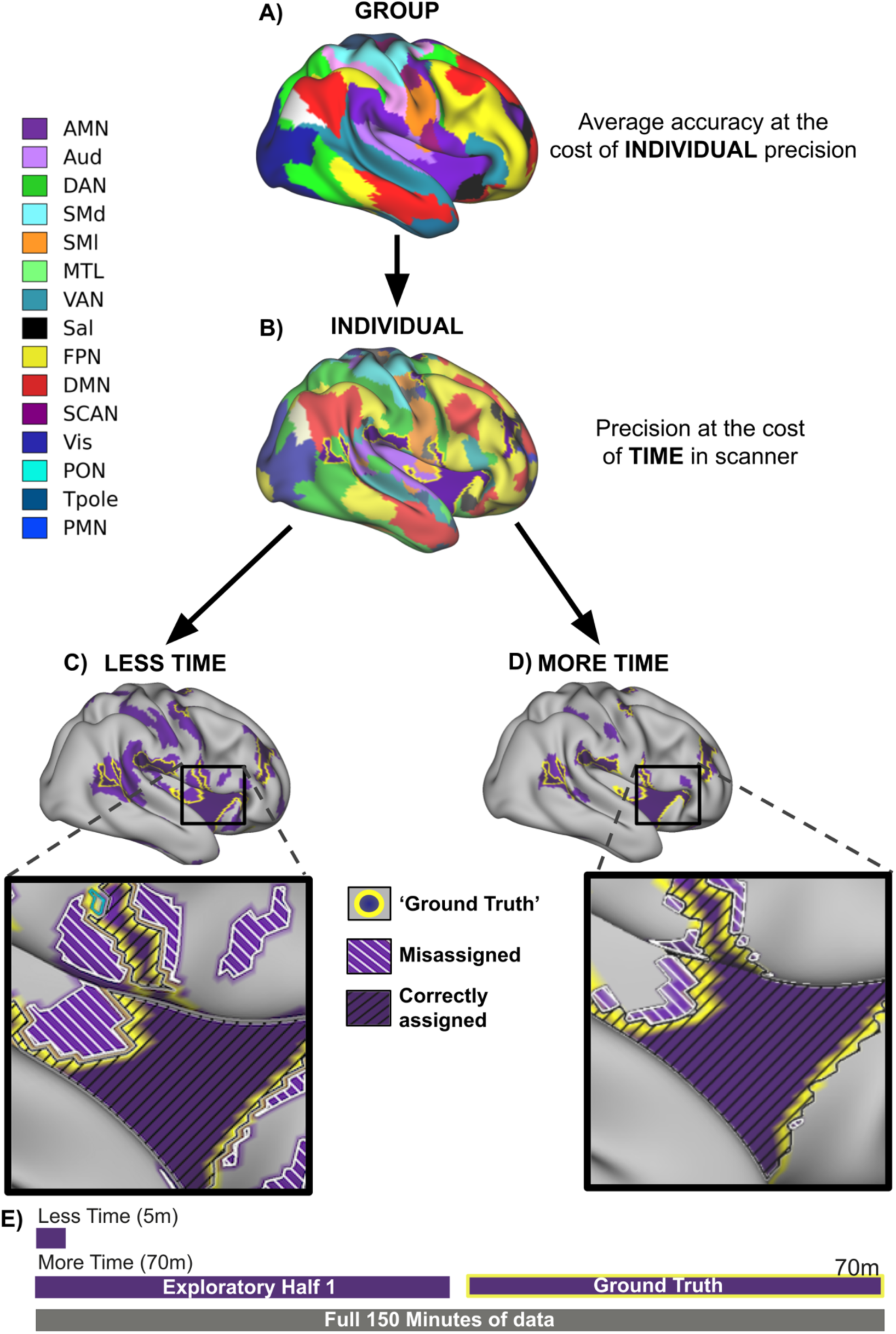
Individualized functional network mapping preserves person-specific topography, but assignment precision depends on data quantity. **A)** Group-average functional network template derived from 141 participants in the Adolescent Brain Cognitive Development (ABCD) study, illustrating the canonical organization of 15 functional networks. Group-average maps provide consistent accurate population-level network definitions but do not capture individual-specific variation in network topography**. B)** Individual-specific functional network maps can produce accurate precision at the cost of time in the scanner. Functional network map is derived from an independent held-out 70 minute resting-state fMRI segment in a densely sampled participant. The cingulo-opercular network, also referred to here as the action mode network (AMN), is outlined in yellow as an illustrative target network**. C–D)** AMN assignments generated using standard template matching from exploratory half 1 with either 5 minutes of data (C, less time) or 70 minutes of data (D, more time). The purple fill indicates the AMN assignment from the exploratory data, and the yellow outline indicates the AMN boundary from the held-out within-subject reference map seen in B. Enlarged lateral views illustrate assignment accuracy, with black diagonal hatching indicating vertices correctly assigned to the AMN in the exploratory and reference data, and white diagonal hatching indicating vertices assigned to the AMN in the exploratory map but to another network in the reference map. The 5 minute assignment contains more misassigned territory, whereas the 70 minute assignment more closely aligns with the held-out reference. **E)** Schematic of the data partitioning procedure. From the full 150 minutes of data collected for each participant, 5 minute and 70 minute segments were drawn from exploratory half 1, while an independent 70 minute segment from half 2 served as the held-out within-subject reference, referred to as the ground truth for brevity. Network abbreviations: AMN, action mode network; Aud, auditory network; DAN, dorsal attention network; SMd, somatomotor dorsal network; SMl, somatomotor lateral network; MTL, medial temporal network; VAN, ventral attention network; SAL, salience network; FPN, frontoparietal network; DMN, default mode network; SCAN, somato-cognitive action network; Vis, visual network; PON, parieto-occipital network; Tpole, temporal pole network; PMN, parietal memory network.

We quantified labeling precision using positive predictive value (PPV), defined as the proportion of each identified network that lies within the corresponding network in the held-out ground truth map. Put simply, PPV indicates how much of the identified network falls within the intended network rather than neighboring networks. This external validation was possible because our densely sampled dataset provided an independent reference half. In a typical dataset containing only 5 to 15 minutes of data, no comparable within-subject reference is available, so the accuracy of the resulting network assignment cannot be directly evaluated. We selected PPV as the primary measure because it specifically penalizes off-network territory included in the identified target, a particularly relevant error for neuromodulation.

As anticipated, PFM-length data (70 minutes of quality data) demonstrated higher PPV than traditional scan-length data (5 minutes of quality data). For example, examining the action mode network (AMN; also known as the cingulo-opercular network; ^29^; Fig. 1C,D), PFM had a PPV of 0.71 ± 0.13 compared to 0.47 ± 0.14 for traditional scan lengths, highlighting the improved alignment with the ground truth reference and reduced misalignment with neighboring networks. Notably, even with longer sampling, the exploratory map did not fully overlap with the 70 minute ground truth reference, which could reflect boundary ambiguity, limited reference duration, or both (Figure 1). This residual ambiguity highlights the need for an internal estimate of assignment stability that can support prioritization of high-confidence network locations when no independent within-subject reference is available, as is typical in short-scan datasets.

### Precision Confidence Mapping (PCM) improves network assignment confidence

To address the limitations of conventional hard-label community detection, we developed Precision Confidence Mapping (PCM), a bootstrap-based framework that quantifies the reliability of individualized network assignments derived from resting-state fMRI data. PCM generates an ensemble of bootstrapped time series datasets and applies community detection independently to each one, producing repeated estimates of network organization within an individual. In this study, PCM was implemented using template matching ^30^, although the framework is compatible with other community-detection approaches such as Infomap ^44^. The principal output is a vertex-wise confidence map that quantifies, for each network, the proportion of bootstrap resamples in which a given vertex was assigned to that network (Fig. 2). These confidence estimates provide a principled way to identify and exclude low-stability vertices prior to neuromodulation targeting.

**Figure 2:**
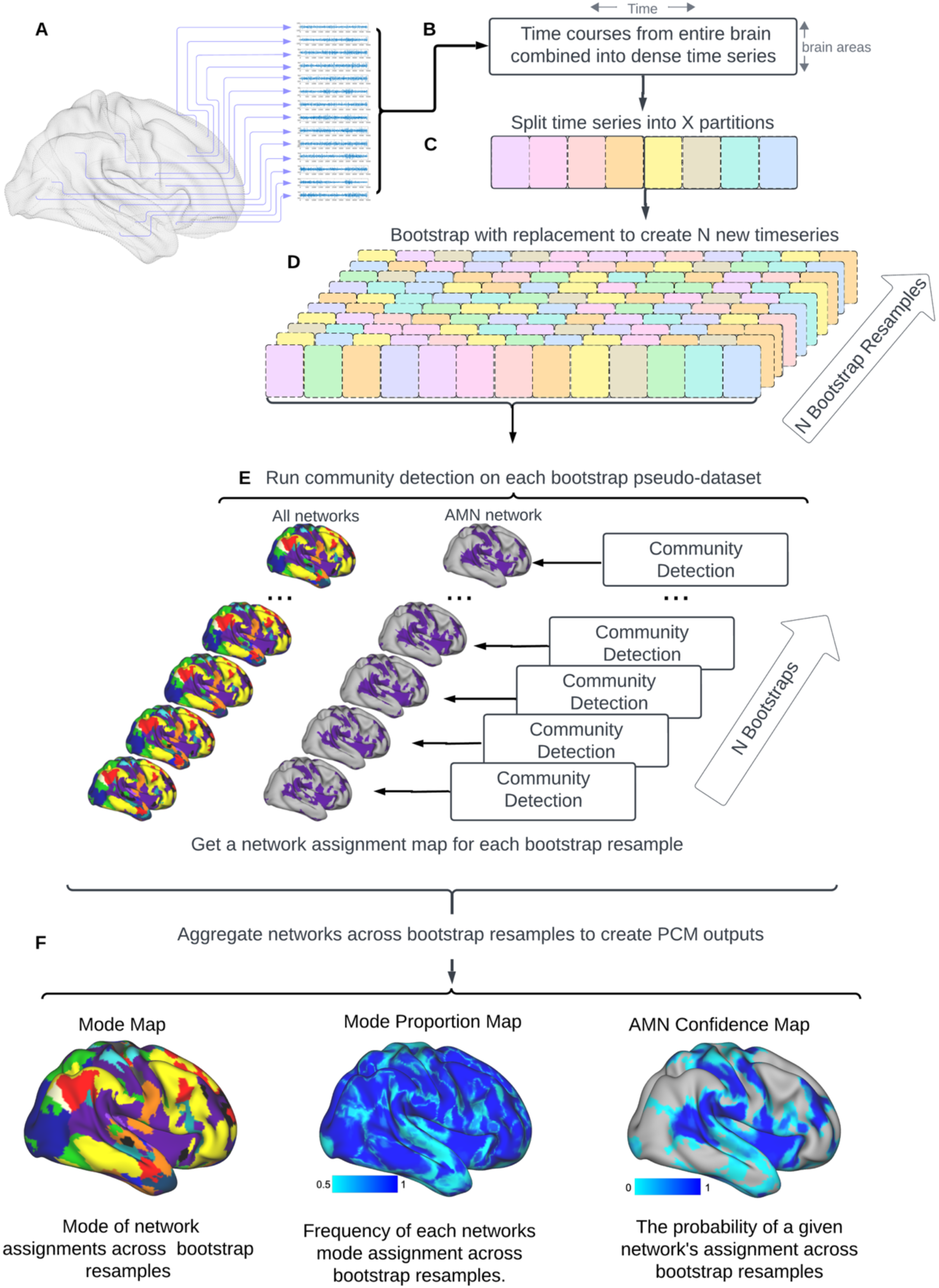
Schematic of precision confidence mapping (PCM) method. **A)** Blood oxygen level-dependent (BOLD) signal fluctuations from each grayordinate across time are combined into a dense time series. **B)** Dense time series comprise BOLD signals from approximately 91,000 grayordinates (y-axis) recorded for each repetition time (TR) across scanning sessions (x-axis). For analyses across scan durations, the dense time series was first restricted to the specified duration, such as 5 or 70 minutes, before PCM was applied. **C)** The selected duration-specific time series was partitioned into resampling units; in the present study, each TR served as one unit. **D)** Resampling with replacement was performed only within the selected time series segment to generate 100 bootstrap pseudo-datasets, each matched in length to the original duration-specific segment. Thus, the 5 minute PCM analysis resampled only the selected 5 minutes of data and did not draw TRs from the larger 70 minute exploratory dataset. **E)** A community detection algorithm (Template Matching) is applied independently to each resampled dataset, resulting in 100 distinct network assignment maps. **F)** Bootstrap resampled results are aggregated to create three output maps: (1) a categorical mode map, assigning each grayordinate to its most frequently occurring network across bootstrap resamples; (2) a mode proportion map, illustrating the frequency of all network assignments across bootstrap resamples; and (3) confidence maps, depicting the probability of assignment to specific networks for each grayordinate.

PCM confidence and PPV quantify distinct properties. Confidence is derived solely from repeated resampling of the available exploratory data, whereas PPV is calculated afterward by comparing the resulting assignments with the independent held-out reference. For each scan duration, bootstrap resampling was restricted to that duration-specific segment. Thus, the 5 minute confidence map was generated entirely from the same 5 minutes of data.

### Thresholding improves AMN targeting

To illustrate how confidence thresholding affected held-out agreement, we applied increasingly stringent confidence thresholds to AMN confidence maps from a representative subject using 5 and 70 minute exploratory scans and compared the retained assignments with the independent 70 minute ground truth reference (Fig. 3). Confidence was defined as the percentage of bootstrap resamples in which each vertex was assigned to the AMN; neither PPV nor the held-out reference was used to determine these confidence values or select vertices for retention. Some correspondence between confidence and PPV may be expected, given that both are based on network assignments generated within the same template-matching framework. This example therefore characterizes how assignment stability relates to held-out reproducibility within that framework, rather than independently validating network identity. In the unthresholded 5 minute map, the AMN extended beyond the ground truth boundary into adjacent regions, indicating substantial false-positive inclusion. As the confidence threshold increased, fewer vertices were retained, and the remaining assignments showed progressively higher agreement with the held-out reference. In the 5 minute scan, PPV increased from 0.35 with no threshold to 0.39 at a 60% confidence threshold, 0.49 at a 90% threshold, and 0.57 at a 99% threshold. By contrast, the 70 minute confidence map showed better agreement with the ground truth before thresholding (PPV = 0.60), with thresholding producing a more modest increase in PPV to 0.70 at the 99% confidence threshold (Fig. 3). This example illustrates two key points. First, short-duration scans contain more low-confidence assignments that can contribute to off-target inclusion. Second, confidence thresholding preferentially removes lower-confidence assignments, improving precision by reducing false-positive inclusion when the goal is network-specific targeting. This framing does not imply that network-border or integrative vertices are universally undesirable, because approaches designed to engage multiple networks may prioritize different target features. We next quantified these effects across all subjects, scan durations, and confidence thresholds.

**Figure 3:**
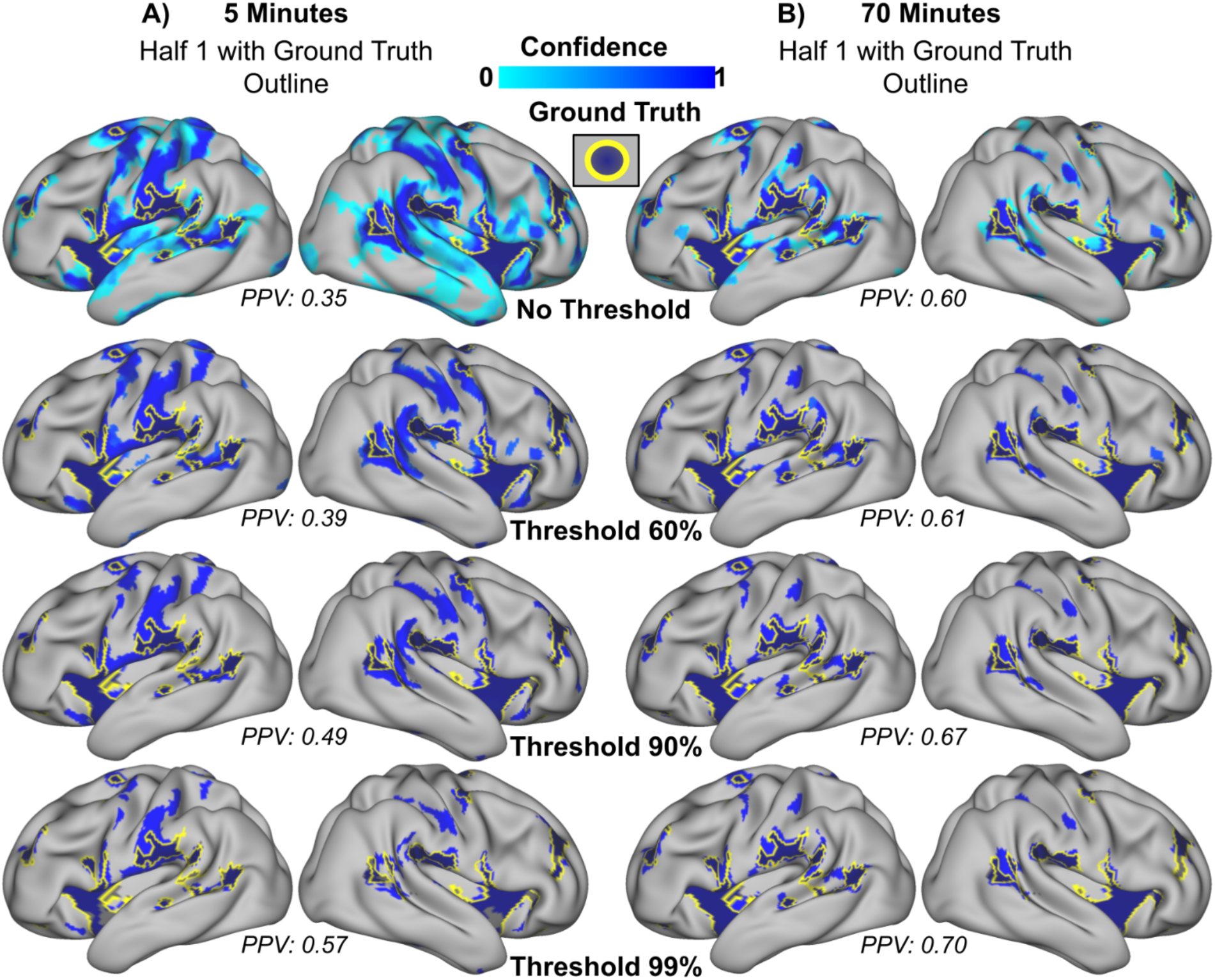
Thresholding confidence maps enhances functional network assignment accuracy by reducing false-positive errors. The action mode network (AMN) is presented as an illustrative example of thresholding effects using different amounts of data. **A)** AMN confidence map derived from 5 minutes of exploratory data (half 1), overlaid with the ground truth AMN network (from 70 minutes of half 2 data, yellow outline). **B)** AMN confidence map derived from 70 minutes of exploratory data (half 1), overlaid with the same ground truth AMN network (yellow outline). For both panels **A** and **B**, the top row depicts unthresholded confidence maps. Subsequent rows display confidence maps thresholded at values of 0.6, 0.9, and 0.99, respectively. Positive Predictive Values (PPV) [True Positives divided by the sum of True Positives and False Positives], are indicated beneath each brain image. Higher thresholds correspond to increased PPV, improving the alignment of the shorter data (5 minute) confidence maps with the ground truth network assignment observed with longer data (70 minute).

### Longer scan durations and confidence thresholding improve AMN targeting precision

We first quantified the effects of scan duration and confidence threshold in the AMN, which served as the primary illustrative network in the main figures. Corresponding analyses across four additional clinically relevant networks and averaged across all 15 canonical networks are presented in the following section and Supplemental Materials. PPV was computed across duration-specific exploratory segments ranging from 5 to 70 minutes and confidence thresholds ranging from 0 to 0.99 (Fig. 4A). For clarity, we highlight representative durations of 5, 25, and 70 minutes of data before framewise displacement censoring and the contrast between unthresholded maps and stringent thresholding at 0.99 in the main text and Table 1, while the full grid of durations, thresholds, and metrics is provided in the Supplemental Materials. Consistent with the illustrative example, PPV increased with both longer exploratory scan duration and more stringent confidence thresholding. For unthresholded maps, PPV improved from 0.47 ± 0.14 at 5 minutes to 0.62 ± 0.15 at 25 minutes and 0.71 ± 0.13 at 70 minutes. Applying a 0.99 confidence threshold further increased PPV at each duration, with the largest gain observed at 5 minutes, where PPV rose from 0.47 ± 0.14 to 0.66 ± 0.14 (Δ = 0.18, t(3) = 15.55, p = 0.0006). At 25 minutes, PPV increased from 0.62 ± 0.15 to 0.74 ± 0.14 (Δ = 0.13, t(3) = 8.80, p = 0.003), and at 70 minutes it increased from 0.71 ± 0.13 to 0.79 ± 0.12 (Δ = 0.08, t(3) = 9.96, p = 0.0022). Thresholding improved PPV at all three representative scan durations, with gains of 0.18, 0.13, and 0.08 at 5, 25, and 70 minutes, respectively. The gain was significantly larger for the shorter 5 minute scans than for the longer 70 minute scans (mean paired difference in gains = 0.103 ± 0.013, t(3) = 16.35, p = 4.98 × 10⁻⁴).

**Figure 4.**
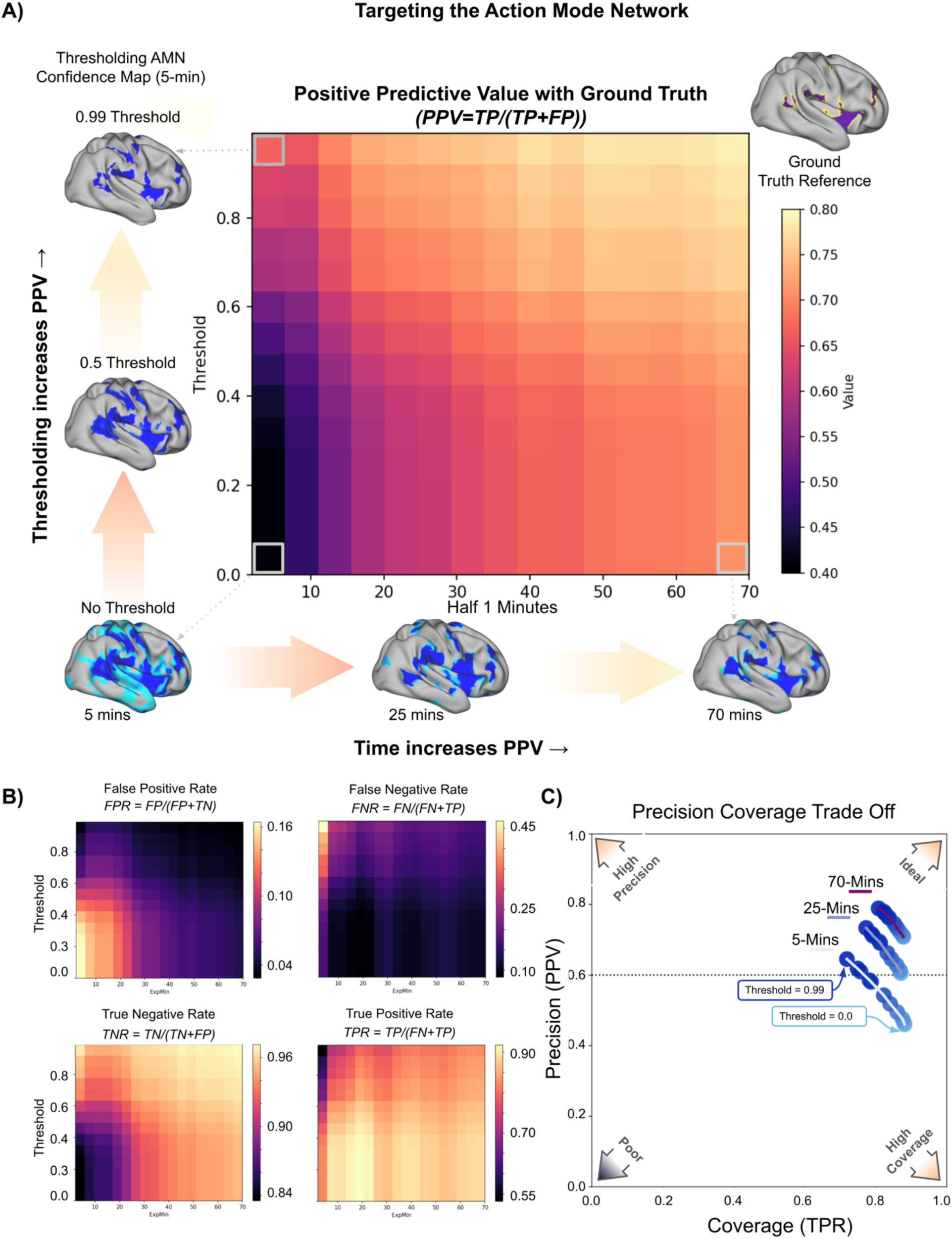
Confidence thresholding and increasing data length improve action mode network (AMN) targeting precision. **A)** Heatmap depicting Positive Predictive Value (PPV; TP / [TP + FP]) as a function of exploratory data length (x-axis) and confidence map thresholding level (y-axis) for the AMN network. Representative confidence maps are shown alongside the axes to illustrate how both data length (e.g., 5–70 minutes) and thresholding (bottom to top) affect the retained network. Heatmap color indicates PPV calculated by comparing network assignments from exploratory half 1 with the ground truth reference derived from half 2 (top right corner). PPV increases with longer data acquisitions and with higher confidence thresholds, suggesting that even short acquisitions can yield high targeting precision when thresholded. Gray boxes and dotted arrows identify the illustrative confidence maps corresponding to selected PPV values. **B)** Complementary heatmaps show additional network comparison metrics. Top left: False Positive Rate (FPR = FP / [FP + TN]) decreases with both thresholding and data length. Bottom left: True Negative Rate (TNR = TN / [TN + FP]) increases correspondingly. Top right: False Negative Rate (FNR = FN / [FN + TP]) increases slightly with high thresholds and short data lengths. Bottom right: True Positive Rate (TPR = TP / [TP + FN]) shows a mild decrease with thresholding but increases overall with data length. **C)** Precision–TPR plot summarizes the tradeoff between targeting precision (PPV, y-axis) and reference-network coverage (TPR, x-axis). Line color reflects data length (minutes), and bubble color indicates threshold level. Longer data generally produce higher PPV and TPR, whereas thresholding increases PPV, particularly at shorter data lengths, at a modest cost to TPR.

**Table 1.** Effects of confidence thresholding on AMN targeting metrics at 5, 25, and 70 minutes.

| Duration | Metric | Thr = 0 | Thr = 0.99 | $\Delta$ | t(3) | p |
| --- | --- | --- | --- | --- | --- | --- |
| **5 minutes** | PPV | $0.47 \pm 0.14$ | $0.66 \pm 0.14$ | 0.18 | 15.55 | 0.0006 |
| | FPR | $0.14 \pm 0.05$ | $0.05 \pm 0.02$ | -0.09 | -6.46 | 0.008 |
| | TNR | $0.86 \pm 0.05$ | $0.95 \pm 0.02$ | 0.09 | 6.46 | 0.008 |
| | FNR | $0.10 \pm 0.08$ | $0.26 \pm 0.12$ | 0.16 | 5.93 | 0.010 |
| | TPR | $0.90 \pm 0.08$ | $0.74 \pm 0.12$ | -0.16 | -5.93 | 0.010 |
| **25 minutes** | PPV | $0.62 \pm 0.15$ | $0.74 \pm 0.14$ | 0.13 | 8.80 | 0.003 |
| | FPR | $0.08 \pm 0.03$ | $0.04 \pm 0.02$ | -0.04 | -5.86 | 0.010 |
| | TNR | $0.92 \pm 0.03$ | $0.96 \pm 0.02$ | 0.04 | 5.86 | 0.010 |
| | FNR | $0.11 \pm 0.09$ | $0.20 \pm 0.13$ | 0.09 | 4.29 | 0.023 |
| | TPR | $0.89 \pm 0.09$ | $0.80 \pm 0.13$ | -0.09 | -4.29 | 0.023 |
| **70 minutes** | PPV | $0.71 \pm 0.13$ | $0.79 \pm 0.12$ | 0.08 | 9.96 | 0.0022 |
| | FPR | $0.05 \pm 0.02$ | $0.03 \pm 0.02$ | -0.02 | -7.34 | 0.005 |
| | TNR | $0.95 \pm 0.02$ | $0.97 \pm 0.02$ | 0.02 | 7.34 | 0.005 |
| | FNR | $0.11 \pm 0.07$ | $0.17 \pm 0.09$ | 0.07 | 5.99 | 0.009 |
| | TPR | $0.89 \pm 0.07$ | $0.83 \pm 0.09$ | -0.07 | -5.99 | 0.009 |

**Note.** Values are mean ± SD across subjects. Thr = 0 and Thr = 0.99 correspond to unthresholded and thresholded maps, respectively. $\Delta$ values were computed as threshold 0.99 minus threshold 0. Paired two-sided t tests were performed across subjects. PPV was the primary targeting metric. FPR and TNR index off-target exclusion, whereas TPR and FNR index the retention and omission, respectively, of reference-defined target territory. AMN denotes the action mode network.

Complementary targeting metrics showed the expected precision-coverage tradeoff (Table 1; Fig. 4B). Thresholding reduced false positive rate (FPR) and increased true negative rate (TNR) at all scan durations, indicating improved exclusion of off-network vertices. These gains were accompanied by lower true positive rate (TPR) and higher false negative rate (FNR), indicating reduced coverage of the reference-defined network. However, the reduction in false positives outweighed these TPR costs for precision targeting.

Confidence thresholding allowed short scans to approach the precision obtained from substantially longer unthresholded scans. At 5 minutes, PPV reached 0.66 ± 0.14 with a 0.99 confidence threshold, compared with 0.71 ± 0.13 using 70 minutes without thresholding. The precision-TPR plot further illustrates this tradeoff (Fig. 4C). Longer scan durations generally improved both PPV and TPR, whereas thresholding shifted performance toward higher PPV at the cost of lower TPR. Together, these results show that PCM thresholding provides a practical route to higher-precision network targeting from short, clinically realistic scan durations.

### Generalizability across clinically relevant networks

To assess whether the effects observed in the AMN generalized beyond a single target network, we performed analogous supplemental analyses in four additional clinically relevant networks: default mode (DMN), frontoparietal (FPN), somato-cognitive action (SCAN), and salience (SAL) (Supplemental Fig. 1). Across all four networks, confidence thresholding produced the same overall pattern observed in the AMN: PPV increased, particularly at shorter scan durations, while true positive rate (TPR), reflecting reference-network coverage, decreased modestly. The DMN showed the highest baseline PPV and TPR, whereas the smaller FPN, SCAN, and SAL networks showed somewhat larger reductions in TPR with thresholding, potentially reflecting their smaller spatial extent and the greater proportion of network territory near boundaries. Averaged across all 15 canonical networks, PPV increased with confidence thresholding at each scan duration, from 0.67 ± 0.04 to 0.86 ± 0.03 at 5 minutes, from 0.71 ± 0.04 to 0.85 ± 0.02 at 25 minutes, and from 0.75 ± 0.03 to 0.85 ± 0.02 at 70 minutes. These increases were significant at all three durations (5 minute: Δ = 0.19, t(3) = 7.87, p = 0.004; 25 minute: Δ = 0.14, t(3) = 22.51, p < 0.001; 70 minute: Δ = 0.10, t(3) = 14.15, p < 0.001), supporting PCM thresholding as a general strategy for improving targeting precision beyond the AMN (Supplemental Fig. 2). Complete descriptive and inferential statistics for each network, threshold, and scan duration are provided in Supplemental Figs. 1–2 and Supplemental Table 1.

### PCM improves short-scan agreement without reducing subject specificity

The preceding analyses showed that confidence thresholding increased AMN targeting precision within participants (Fig. 4). However, this improvement could reflect either preferential retention of stable, participant-specific AMN territory or convergence on a generic AMN core shared across participants. To distinguish these possibilities, we compared each participant’s experimental AMN map with both their own fixed 70 minute reference, reflecting within-subject agreement, and the references of the other three participants, reflecting between-subject agreement. If PCM preserved subject specificity, it should increase within-subject agreement without reducing the advantage of within-subject over between-subject agreement.

For each participant, AMN maps were generated from 5 or 70 minutes of exploratory data using standard template matching (TM) or PCM confidence maps thresholded at 0.99. TM served as the unthresholded baseline, and each 5 minute PCM map was generated solely from its selected 5 minute segment. Within-subject PPV was calculated against the same participant’s fixed, unthresholded 70 minute reference. Between-subject PPV was calculated as the mean agreement with the other three participants’ references, yielding one between-subject value per participant and condition (Fig. 5). The same fixed reference maps were used throughout all comparisons.

**Figure 5.**
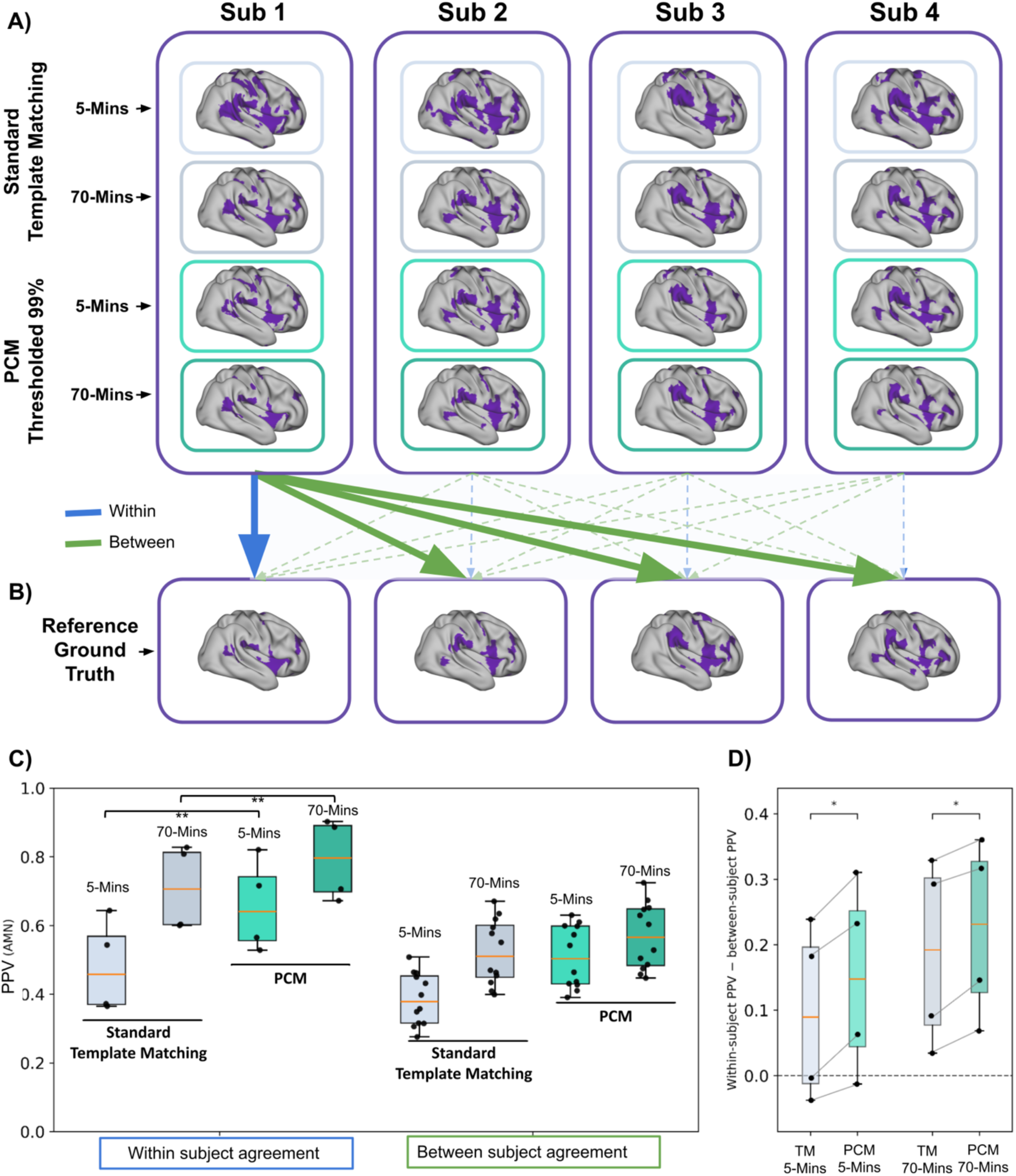
PCM improves short-scan agreement while preserving subject-specific AMN topography. **A)** AMN network maps for each participant (Sub1 through Sub4) estimated from short (5 minute) and long (70 minute) data using standard template matching (top two rows) or PCM confidence maps thresholded at 0.99 (bottom two rows). Purple indicates vertices assigned to the AMN. **B)** Fixed subject-specific reference, or ground truth, AMN maps derived from each participant’s 70 minute reference data. Arrows illustrate the comparison structure. Within-subject comparisons, shown in blue, match each experimental map with the same participant’s 70 minute reference. Between-subject comparisons, shown in green, match each experimental map with the 70 minute references of the other participants. **C)** Positive predictive value (PPV) of AMN assignments relative to the fixed 70 minute references. The left group shows within-subject PPV, with one value per participant, for TM and PCM at 5 and 70 minutes. The right group displays all 12 between-subject pairwise comparisons for visualization. For inferential analyses, the three between-subject comparisons were averaged within participant so that each participant contributed one value per condition. Boxplots show the median and interquartile range, whiskers extend to the most extreme non-outlier values, and points show individual comparisons. **D)** Subject-specificity separation, defined as within-subject PPV minus mean between-subject PPV, for each participant and condition. Positive values indicate greater agreement with a participant’s own 70 minute reference than with the other participants’ references. Each point represents one participant, and thin lines connect the same participant’s TM and PCM values at matched scan duration. The dashed line marks zero separation. Holm-corrected paired comparisons showed higher within-subject PPV for PCM than TM at both 5 and 70 minutes (p Holm < 0.01) and greater within-versus-between separation for PCM at both durations (p Holm = 0.025).

PCM increased agreement with each participant’s own reference at both scan durations. At 5 minutes, within-subject PPV increased from 0.48 ± 0.14 with TM to 0.66 ± 0.14 with PCM (Δ = 0.177 ± 0.018, paired t(3) = 19.76, p = 2.83 × 10⁻⁴, p_Holm = 0.00113). At 70 minutes, within-subject PPV increased from 0.71 ± 0.12 to 0.79 ± 0.12 (Δ = 0.082 ± 0.016, t(3) = 10.02, p = 0.00212, p_Holm = 0.00635). The larger numerical gain was observed for the 5 minute maps.

We next assessed whether this improvement came at the cost of subject specificity. For each participant and condition, we calculated a subject-specificity separation score, defined as within-subject PPV minus mean between-subject PPV. Positive values indicate that an experimental map more closely resembles the participant’s own reference than the references of other participants. If PCM homogenized maps toward a shared AMN core, this separation would be expected to decrease. Mean separation was positive in all four conditions: 0.095 ± 0.136 for TM at 5 minutes, 0.148 ± 0.149 for PCM at 5 minutes, 0.187 ± 0.146 for TM at 70 minutes, and 0.223 ± 0.138 for PCM at 70 minutes (Fig. 5D). Rather than decreasing, separation increased under PCM at both 5 minutes (Δ = 0.053 ± 0.021, t(3) = 5.06, p = 0.0149, p_Holm = 0.0245) and 70 minutes (Δ = 0.036 ± 0.013, t(3) = 5.43, p = 0.0122, p_Holm = 0.0245; Fig. 5D). Together, these results indicate that PCM improved agreement with each participant’s own network topography without reducing subject specificity in this densely sampled cohort.

## Discussion

Precise neuromodulation depends on accurately identifying the functional network one intends to stimulate, but most individualized resting-state mapping approaches return hard network labels without indicating how stable those assignments are to variation in the available data^25,30,47^. In this study, we addressed that gap with Precision Confidence Mapping, a bootstrap-based framework that estimates per-vertex assignment confidence and enables low-stability regions to be excluded before targeting. Across scan durations, PCM thresholding increased targeting precision, with the largest gains observed in short scans where standard community detection methods performed least well. Thresholding reduced FPR and increased TNR with only modest costs to TPR, generalized across additional clinically relevant cortical networks, and improved agreement with a fixed 70 minute reference while preserving the separation between within-subject and between-subject agreement. Together, these findings suggest that PCM provides a practical route to more cautious and precise individualized network targeting using scan durations commonly acquired in clinical and translational studies.

These findings are especially relevant in clinical and translational settings, where individualized resting-state mapping is typically performed with scan durations far shorter than those used in dense-sampling studies. Clinical resting-state acquisitions often last only 5 to 15 minutes, whereas stable individualized network estimates can require substantially more data, in some cases exceeding 100 minutes ^14,24,39–42^. When acquisitions are this brief, individualized targets may still be derived, but the reliability of the resulting map is often unknown.

Functional connectivity-guided targeting has shown clear promise in experimental, mechanistic, and some individualized treatment studies, where accounting for subject-specific network anatomy can improve engagement of the intended circuit ^9,34,48^. However, evidence from clinical trials in major depressive disorder has been less consistent, and recent reviews have questioned whether individualized imaging-guided targeting is superior to standard approaches across studies ^49^. One plausible contributor to this discrepancy is that clinical imaging often relies on relatively short resting-state scans or simplified approaches that provide limited information about full network topography and no direct estimate of map reliability.

PCM is designed to address this specific problem by adding a reliability layer to individualized network maps. Rather than replacing community-detection methods, PCM builds on them by estimating per-vertex assignment confidence across repeated resamples. PCM confidence should not be interpreted as a direct measure of signal-to-noise ratio. Signal-to-noise measures characterize properties of the underlying BOLD signal, whereas PCM quantifies the stability of the resulting network assignment after the full mapping procedure is repeated across bootstrap samples. Low signal quality may contribute to low confidence, but assignment instability may also arise when a vertex’s connectivity profile is similarly compatible with multiple networks or when its label is sensitive to the network-identification procedure.

This assignment-level information is especially useful where targeting decisions are hardest, namely along network borders and in short, noisy scans, where standard hard-label approaches typically treat all assigned vertices as equally valid. In our data, standard template matching from short scans produced substantial off-network contamination, whereas PCM thresholding recovered much of that lost precision. As illustrated in our conceptual framework (Fig. 6), this distinction matters because, for neuromodulation, all assignment errors are not equally consequential. From a clinical targeting perspective, an optimal target should reliably engage the intended network while minimizing inadvertent stimulation of adjacent, functionally distinct territories ^23,50,51^. Even when the intended network is identified broadly, unstable border regions may still be incorporated into the target as though they were as reliable as the network core. For this reason, we emphasize positive predictive value as a clinically relevant measure of targeting precision. In this framing, a modest reduction in TPR may be acceptable so long as enough true positive vertices remain for stimulation, particularly if that tradeoff substantially reduces off-target inclusion and leaves a target composed of the vertices most confidently assigned to the intended network.

**Figure 6:**
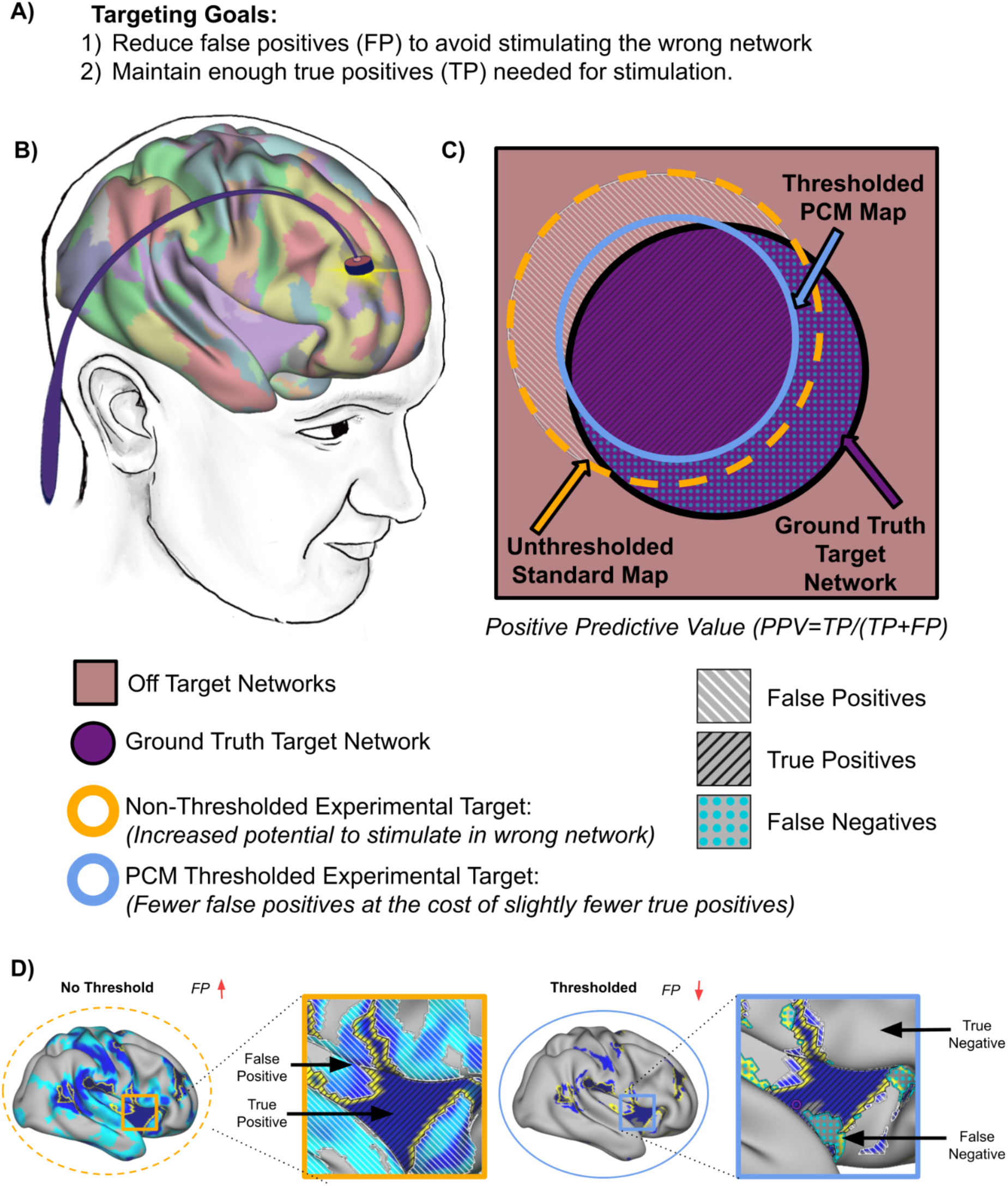
Schematic illustrating key considerations in targeting specific brain areas. **A)** Emphasizes the importance of minimizing false-positive (FP) assignments while retaining a sufficient number of true-positive (TP) assignments to accurately stimulate the intended target network. **B)** Cartoon depiction of how multiple functional networks are spatially intermingled across the cortical surface, highlighting the complexity of accurate network targeting. **C)** Cartoon example illustrating risks of excessive FPs. The solid purple circle marks the intended ground truth target area. The dark yellow dotted outline represents an unthresholded experimental target derived from standard community detection, showing considerable overlap with hypothetical off target networks. The solid light-blue outline demonstrates how applying confidence thresholding shrinks this experimental target, reducing FPs while preserving adequate number of TPs. Classification outcomes are marked using the same hatch/dot convention as the brain surfaces in panel D: light diagonal hatching indicates FPs (off-target areas covered by either experimental map, but outside of the ground truth network), dark diagonal hatching indicates TPs (on-target areas covered by both experimental map and ground truth network), and light dotted fill indicates FNs (ground truth areas missed by the thresholded target). The light red square surrounding the target represents off-target networks that could be inadvertently stimulated due to FPs. Positive Predictive Value (PPV), a measure of precision, is calculated as TP / (TP + FP). **D)** Illustrates this concept using an AMN (action mode network) confidence map. A comparison between a non-thresholded and a thresholded confidence map is shown, with ground truth overlaid. The example compares the experimental half1-derived network assignment to a 70 minute ground truth. The brain schematic demonstrates that thresholding reduces FPs, though at a minor cost of reducing TPs.

A related concern is that thresholding may improve precision by discarding the very portions of a map that contribute to an individual’s unique network organization. That possibility is not trivial, because subject-specific functional topography can itself be clinically meaningful, including in depression, where large-scale network alterations can vary across individuals ^52^. If confidence thresholding merely removed variable regions and left only a generic network core, then gains in apparent reliability would come at the expense of individual specificity. Our within-subject versus across-subject analyses do not support that interpretation. PCM markedly increased short-scan agreement with the fixed 70 minute reference while preserving, and in this dataset even increasing, the separation between within-subject and across-subject agreement. These findings suggest that PCM preserves individualized topography by enriching for vertices that are stably and specifically assigned within each individual.

Importantly, preserving subject-specific topography is only clinically useful if the framework also generalizes beyond a single illustrative network. The AMN served as the primary illustrative example, while the supplemental analyses show that PCM generalizes to other individualized functional networks. These analyses demonstrate average thresholding benefits across all 15 template networks, with more detailed characterization of four clinically relevant networks: SCAN, DMN, FPN, and salience (Supplemental Figs. 1–2). These additional networks were included to test generalizability, and also because contemporary neuromodulation is increasingly motivated by distributed circuit and network models rather than by purely anatomical one-site-fits-all targeting rules ^9,16,21,53^. These networks are clinically relevant because they are altered across patient populations and provide plausible circuit-level substrates for individualized neuromodulation. For example, SCAN has recently been implicated in Parkinson’s disease pathophysiology and treatment response, with successful neuromodulation reducing SCAN hyperconnectivity and SCAN-targeted TMS outperforming effector-based targeting ^54^. SCAN and AMN have also been proposed as candidate precision targets for chronic pain neuromodulation, consistent with evidence that these networks participate in action-relevant pain processing ^28,29,36^. In addition, the salience network shows trait-like topographic expansion in depression ^52^, frontoparietal control systems are implicated across disorders of executive control including depression and ADHD ^55,56^, and the default mode network is altered across conditions including depression and schizophrenia ^57–61^. Viewed in this way, PCM supports target refinement across individualized functional networks by isolating the most stably assigned portions of the network most relevant to a given disorder.

That broader applicability does not eliminate important methodological tradeoffs. One methodological consideration in PCM is the treatment of temporal order. By resampling individual functional volumes, PCM prioritizes stable network topography over moment-to-moment variation, consistent with prior work showing that individualized functional networks are dominated by stable subject-level features ^24,25,32,40^. At the same time, this resampling disrupts the original temporal autocorrelation structure and deemphasizes transient state-dependent fluctuations that are intrinsic to fMRI time series. Prior work suggests that transient fluctuations can reflect biologically meaningful spontaneous co-activation structure and time-varying brain states ^62–65^. More broadly, temporal autocorrelation itself appears to be a reliable and informative property of rs-fMRI data, even though its estimation is also shaped by methodological and statistical constraints ^66^. PCM is therefore better suited for estimating stable network topography than for preserving rapid moment-to-moment changes in brain state. This tradeoff is reasonable for the present goal, because neuromodulation targeting typically depends on identifying reproducible spatial network structure rather than capturing brief dynamic fluctuations.

A second methodological consideration is that PCM confidence estimates still depend, in part, on the underlying network-identification procedure. The choice of template prior, for example, may be especially relevant in developmental or clinical populations, where functional organization can differ from normative adult templates ^25,30^. This limitation is more properly attributed to the base mapping framework than to PCM itself, and, in principle, PCM could be extended across multiple atlases, parameter settings, or network-detection approaches. Similarly, the present results focus on cortical functional networks, in part because subcortical functional organization is harder to map reliably owing to lower signal-to-noise ratio and less mature template resources. However, confidence mapping may prove especially useful in subcortical territories, where imprecision in localization can be particularly consequential. Developing stronger individualized subcortical templates, potentially leveraging ultra-high-field 7T fMRI and related high-resolution mapping approaches, will therefore be an important next step toward more reliable subcortical network identification, on which PCM can then build.

Beyond these methodological considerations, data quality and quantity remain first-order determinants of reliability ^67^. PCM cannot rescue severely compromised data, and long, low-motion acquisitions with high signal-to-noise ratio still provide the strongest basis for individualized mapping. Here, we leveraged a small but deeply sampled cohort, which was well suited for demonstrating the method’s technical behavior under controlled conditions. However, broader generalizability will require replication in larger and more diverse samples, including clinical and developmental cohorts. Future work should also test whether combining PCM with complementary information, such as structural MRI, EEG, or subject-specific electric-field modeling, can further sharpen target localization and improve the interpretability of individualized neuromodulation maps.

Taken together, these findings position PCM as a practical extension of individualized functional mapping that makes such maps more interpretable and actionable. By quantifying assignment confidence, improving targeting precision under realistic scan constraints, and preserving subject-specific topography, PCM provides a concrete methodological step toward more reliable precision neuromodulation.

## Methods

### 1. Resource Availability

#### Lead Contact

Further information and requests for resources and reagents should be directed to and will be fulfilled by the Lead Contact, Julian Sergej Benedikt Ramirez.

#### Materials Availability

This study did not generate new, unique reagents or materials.

#### Data and Code Availability

All data reported in this paper will be shared by the lead contact upon request. The code used for analysis is available upon request to the lead author.

#### Declaration of competing interests

J.S.B.R., R.J.M.H., D.A.F., and S.M.N. are named inventors on a provisional patent application related to Precision Confidence Mapping and the community detection methods described in this manuscript. D.A.F. and N.U.F.D. have a financial interest in Turing Medical Inc. and may benefit financially if the company is successful in marketing FIRMM motion monitoring software products. D.A.F. and N.U.F.D. may receive royalty income based on FIRMM technology developed at Washington University School of Medicine and Oregon Health & Science University and licensed to Turing Medical Inc. D.A.F. and N.U.F.D. are cofounders of Turing Medical Inc. The interests involving Turing Medical have been reviewed and are managed by Washington University School of Medicine, Oregon Health & Science University, and the University of Minnesota. All other authors declare no competing interests.

#### Funding

J.S.B.R. T32 Training Grant: 5T32DA007234-37

B.T.C. K23DA057486

O.M.D. Research reported in this publication was supported by the University of Minnesota’s MnDRIVE (Minnesota’s Discovery, Research and Innovation Economy) initiative.

### 2. Experimental Model and Subject Details

#### Participants

Data were collected from four adults (three females, one male) with a mean age of 29.5 years (SD = 6.56). These data were collected as part of a larger precision functional mapping (PFM) study comparing 3-tesla and 7-tesla resting-state functional magnetic resonance imaging (rsfMRI) scans. Each participant completed three sessions at 3 tesla and three sessions at 7 tesla in interleaved order, with 50 minutes of fixation (cross-hair) resting-state data collected at each session. All sessions for a given participant were completed within 6 weeks (M = 31.75; SD = 6.29 days). For the current study, only 3-tesla scans were utilized. Subjects were enrolled at the University of Minnesota (UMN), with an inclusion criterion of being between 18 and 50 years of age at the time of enrollment. Exclusion criteria included ferromagnetic metallic implants, claustrophobia, or other contraindications to MRI, pregnancy, and non-English speakers. The study was approved by the University of Minnesota’s Institutional Review Board, with written informed consent obtained from participants before the first scan.

### 3. Method Details

#### Data Acquisition

All imaging was acquired on a Siemens 3T MAGNETOM Prisma scanner (Siemens Healthineers, Erlangen, Germany) using a 32-channel receive head coil.

Resting-state fMRI data comprised five 10 minute multiband multi-echo gradient-echo EPI runs per session, with 3 sessions per subject. Each run included 340 volumes acquired with the following parameters: TR = 1761 ms; four echoes (TE = 14.20, 38.94, 63.66, 88.40 ms); 2.0 mm isotropic resolution; 72 axial slices; multiband factor = 6; in-plane GRAPPA acceleration = 2; flip angle = 68°; bandwidth = 2272 Hz/pixel; anterior-to-posterior phase-encoding direction.

Geometric distortions were corrected using paired spin-echo field maps (AP and PA directions) matching the functional scan geometry (72 slices, 2.0 mm isotropic), acquired with parameters: TR = 13693 ms; five echoes (TE = 20.80, 58.48, 96.16, 133.84, 171.52 ms); flip angle = 90°; GRAPPA acceleration = 2; three volumes collected per polarity. An additional 30-volume reverse-polarity multi-echo EPI run with posterior-to-anterior phase encoding was also acquired to further supplement distortion correction.

High-resolution structural imaging consisted of a sagittal T1-weighted MP-RAGE sequence (1.0 mm isotropic resolution; TR = 2500 ms; TE = 2.90 ms; inversion time (TI) = 1070 ms; flip angle = 8°; GRAPPA = 2; bandwidth = 240 Hz/pixel) and a T2-weighted SPACE sequence (1.0 mm isotropic resolution; TR = 3200 ms; TE = 565 ms; GRAPPA = 2; bandwidth = 241 Hz/pixel).

#### Data Preprocessing

Data were first converted into BIDS format using dcm2bids ^68^. Prior to standard preprocessing, thermal noise removal was performed using Noise Reduction with Distribution Corrected (NORDIC) principal component analysis ^69–71^. NORDIC denoising utilized both the phase and magnitude components of the MRI scan. The use of these phase and magnitude data preserves the integrity of complex-valued Gaussian noise characteristics ^70^. Noise estimation leveraged three additional noise volumes acquired at the conclusion of each functional run. NORDIC denoising was conducted in MATLAB R2019a on each echo separately due to the multi-echo (ME) nature of our dataset ^69^.

Following NORDIC denoising, fMRI data underwent preprocessing with fMRIprep (version 24.0.0-dev55, GitHub commit 4d21c37a; ^72^). Fieldmap-based susceptibility distortion correction for the BOLD time series utilized FSL’s “topup” algorithm, estimating the fieldmaps from paired acquisitions with reversed phase-encoding directions (PEPolar method; ^73^). Additional preprocessing parameters included the “--project-goodvoxels” flag, ensuring voxels exhibiting high local variation were omitted during the projection onto the cortical surface, and the “--cifti-output 91k” flag to generate outputs in the HCP grayordinate space ^74^. Echo combination in fMRIprep was optimized via a T2*-weighted approach (wTE = TE * exp(-TE/T2*); ^75^).

Post-processing was carried out using XCP-D version 0.6.1 ^76^. Configurations used included: “--cifti” for inputting fMRIprep-generated CIFTI derivatives; “--warp-surfaces-native2std” to apply non-linear transformations aligning outputs to the MNI152NLin6Asym template; “-m” for concatenating functional and motion time series data across runs; “--dcan-qc” to generate comprehensive quality control reports analogous to DCAN’s ABCD-BIDS pipeline summaries; “--despike” using AFNI’s 3dDespike method; “--motion-filter-type notch” with respiratory filtering set between “--band-stop-min 12” and “--band-stop-max 18,” based on recommended respiratory frequency ranges from XCP-D documentation.

Nuisance regression included 36 parameters: six motion regressors (with temporal derivatives and quadratic terms), average signals from gray matter, white matter, and cerebrospinal fluid (each with temporal derivatives and quadratic terms). Band-stop filtering of motion parameters was applied before regression. Frames exceeding a Framewise Displacement (FD) threshold of 0.3 mm were interpolated using adjacent lower-motion frames prior to denoising. A stricter FD threshold of 0.2 mm was employed when censoring frames for subsequent functional connectivity matrix computations. Prior to analyses, executive summaries from XCP-D and visual quality control files from fMRIprep were reviewed to ensure that data quality met standards.

#### Precision Confidence Mapping (PCM)

Preprocessed BOLD time series from approximately 91k grayordinates in CIFTI format were used for all analyses. Motion scrubbing was performed by excluding frames with framewise displacement (FD) greater than 0.2 mm ^42^.

PCM begins by segmenting the dense time series into temporal units for bootstrap resampling. The method is flexible and allows the user to define the resampling unit as individual TRs, contiguous time blocks of any chosen duration (for example, 1 to several minutes), or proportions of the full dataset divided into larger chunks. By default, however, PCM uses bootstrap resampling with replacement at the level of individual TRs. In the present study, we used this default approach to generate 100 pseudo-datasets, each matched in length to the original scan. Resampling individual TRs provides repeated estimates of network organization within an individual, stabilizing network topography by averaging over moment-to-moment fluctuations, although at the cost of reduced sensitivity to transient state-dependent dynamics ^32^. Each pseudo-dataset then independently undergoes community detection to assign every grayordinate to a functional network (Figure 2).

Although PCM is compatible with various clustering algorithms, this study employed Template Matching (TM), as detailed by ^30^. In brief, TM assigns network labels based on the spatial similarity between individual connectivity profiles and a predefined set of network templates. The template set used here was generated from an independent cohort of 141 ABCD participants (mean age = 9.92 ± 0.63 years; n = 67 female, 47.5%; n = 74 male, 52.5%), using previously established adult network priors from the WashU 120 cohort ^25^. Because the somato-cognitive action network (SCAN) was not included in those original priors, it was incorporated separately based on the SCAN network described ^28^. The resulting template set delineated 15 canonical functional networks: default mode (DMN), visual (Vis), frontoparietal (FPN), dorsal attention (DAN), ventral attention (VAN), salience (SAL), action mode network (formerly cingulo-opercular), somatomotor dorsal (SMd), somatomotor lateral (SMl), auditory (Aud), temporal pole (Tpole), medial temporal lobe (MTL), parieto-occipital (PON), parietal memory (PMN), and somato-cognitive action (SCAN) networks.

To incorporate SCAN into the template matching process, we adapted the procedure by initially computing seed-based correlations using an average BOLD time series across grayordinates within an independent fMRI dataset. These correlation values were averaged across participants, generating network-specific correlation maps for each grayordinate. Each network template underwent thresholding at correlation Z-scores greater than 1 (top ∼15.9% of connections). Grayordinate assignments to networks were based on calculating eta² values, a measure of map-to-map similarity, selecting the network with the highest eta² value. For motor networks specifically, a higher correlation threshold (Z-score > 3) was applied to differentiate SCAN from other motor-related networks.

The stability of the network assignments derived from bootstrapped resamples was quantified by collapsing the results into three complementary brain maps. First, a mode map was produced, assigning each grayordinate the network label most frequently occurring across the 100 bootstrap resamples. Second, a mode-proportion map was generated to quantify the consistency of the modal label by depicting the fraction of resamples assigning the grayordinate to its mode network. Lastly, voxel-wise confidence maps were calculated for each network by determining the proportion of bootstrap resamples in which each grayordinate was assigned to each network. Collectively, these maps offer categorical (mode) and probabilistic (mode-proportion and confidence maps) assessments of assignment reliability. This aggregation approach parallels the probabilistic atlas methodology described by ^30^ but is uniquely applied across multiple bootstrap resamples within an individual, thereby enhancing the precision and reliability of personalized network identification.

#### Reliability Testing

To assess the reliability of PCM and evaluate whether confidence maps effectively identify less reliable network assignments arising from shorter data segments, we employed a split-half analysis strategy using our extensive (∼150 minute) dataset. The dataset was divided into two halves: an exploratory half (half 1) and a ground truth half (half 2). The exploratory half was further segmented into incremental bins of varying durations (5, 10, 15, 20, 25, 30, 35, 40, 45, 50, 55, 60, 65, and 70 minutes). PCM and standard template matching were applied independently to these incremental bins as well as to the full 70 minute ground truth reference half.

Network assignments from each incremental exploratory bin were directly compared to assignments from the 70 minute ground truth reference. Precision and reliability were quantified using several metrics, including the Positive Predictive Value (PPV), defined as: *PPV* = *TP*/(*TP* + *FP*) where *TP* represents true positives, and *FP* represents false positives. Additional metrics evaluated included the True Positive Rate (TPR; sensitivity): *TPR* = *TP*/(*TP* + *FN*); True Negative Rate (TNR; specificity): *TNR* = *TN*/(*TN* + *FP*); False Positive Rate (FPR): *FPR* = *FP*/(*FP* + *TN*); and False Negative Rate (FNR): *FNR* = *FN*/(*FN* + *TP*). Normalized Mutual Information (NMI) was also computed to assess overall agreement between exploratory and ground truth network assignments: 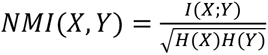 where *I(X;Y)* represents mutual information, and *H(X)* and *H(Y)* denote the entropies of the exploratory and ground truth assignments, respectively.

To further evaluate the precision improvements achievable through PCM, confidence maps derived from each incremental bin were thresholded at multiple confidence levels (0, 0.1, 0.2, 0.3, 0.4, 0.5, 0.6, 0.7, 0.8, 0.9, and 0.99). At each threshold, network assignments from exploratory bins were compared to the stable assignments obtained from the 70 minute ground truth reference using standard template matching. These comparisons aimed to identify threshold levels that significantly enhanced precision, thereby informing optimal thresholding practices for reliable network targeting.

The effectiveness of confidence map thresholding and resulting improvements in precision were visualized and evaluated through heatmaps and precision-TPR curves, facilitating the identification of data length and thresholds yielding stable and reliable network assignments.

#### Statistical Analyses

##### Thresholding effects on performance metrics (Table 1; Fig. 4)

To quantify how confidence thresholding changed AMN network performance, we extracted subject-level values for cortical metrics (PPV, TPR, TNR, FPR, FNR) at three scan lengths (5, 25, and 70 minutes), evaluated against a fixed 70 minute reference. We focused inferential tests on two threshold conditions: unthresholded maps (threshold = 0) and high-confidence maps (threshold = 0.99). Metrics were paired within subjects (N = 4). We report mean ± SD across subjects for each Metric × Scan-length × Threshold cell. To test whether thresholding altered each metric at a given scan length, we used two-tailed paired t-tests comparing threshold 0 vs 0.99. Because the sample size was small and metrics are bounded in [0,1], we also ran an exact Wilcoxon signed-rank test as a nonparametric robustness check; conclusions were qualitatively consistent. Effect sizes were quantified using the paired-sample, small-sample bias–corrected Hedges’ g. For these scan-length comparisons, p-values are reported unadjusted and interpreted descriptively.

Within- and between-subject comparisons against a fixed reference (Fig. 5): AMN maps were generated for each participant using standard template matching (TM; unthresholded) or PCM confidence maps thresholded at 0.99, applied to 5- or 70 minute exploratory segments, and compared against that participant’s fixed, unthresholded 70 minute reference map. Within-subject PPV was defined as the PPV between each participant’s experimental map and their own reference (Exp(i) vs. Ref(i); diagonal comparisons). Between-subject PPV was computed for the three off-diagonal comparisons to the other participants’ references (Exp(i) vs. Ref(j≠i)) and averaged within participant before any statistical comparison, to avoid pseudoreplication. Subject-specificity separation was defined as within-subject PPV minus mean between-subject PPV. We report mean ± SD for within-subject PPV, between-subject PPV, and separation for each condition (TM 5 minute, TM 70 minute, PCM 5 minute, PCM 70 minute). Inferential testing was limited to a pre-specified set of four paired comparisons across subjects (N = 4): within-subject PPV differences between PCM and TM at matched duration (5 minute and 70 minute), and differences in within–between separation between PCM and TM at matched duration (5 minute and 70 minute), using two-tailed paired t-tests with Holm–Bonferroni correction applied across the four comparisons. All analyses were performed in Python using pandas and SciPy with custom code.

## Notes

J.S.B.R: T32 NIDA Training Grant [5T32DA007234-37]; B.T.C. K23DA057486; O.M.D. Research reported in this publication was supported by the University of Minnesota’s MnDRIVE (Minnesota’s Discovery, Research and Innovation Economy) initiative.

## References

1. Hyde, J. et al. Efficacy of neurostimulation across mental disorders: systematic review and meta-analysis of 208 randomized controlled trials. Mol. Psychiatry 27, 2709–2719 (2022).

2. Mehta, D. D. et al. A systematic review and meta-analysis of neuromodulation therapies for substance use disorders. Neuropsychopharmacology 49, 649–680 (2024).

3. Sahay, S. et al. Harnessing neuroimaging-guided transcranial magnetic stimulation for precision therapy in substance use disorders. Mol. Psychiatry 30, 3804–3816 (2025).

4. Krauss, J. K. et al. Technology of deep brain stimulation: current status and future directions. Nat. Rev. Neurol. 17, 75–87 (2021).

5. Dougherty, D. D. Deep brain stimulation: Clinical applications. Psychiatr. Clin. North Am. 41, 385–394 (2018).

6. Nahas, Z., et al. Personalized Adaptive Cortical Electro-stimulation (PACE) in Treatment-Resistant Depression. (2025).

7. Cox, S. S., Connolly, D. J., Peng, X. & Badran, B. W. A comprehensive review of low-intensity focused ultrasound parameters and applications in neurologic and psychiatric disorders. Neuromodulation 28, 1–15 (2025).

8. Boccard, S. G. J., Pereira, E. A. C. & Aziz, T. Z. Deep brain stimulation for chronic pain. J. Clin. Neurosci. 22, 1537–1543 (2015).

9. Cash, R. F. H. & Zalesky, A. Personalized and circuit-based transcranial magnetic stimulation: Evidence, controversies, and opportunities. Biol. Psychiatry 95, 510–522 (2024).

10. Agboada, D., Zhao, Z. & Wischnewski, M. Neuroplastic effects of transcranial alternating current stimulation (tACS): from mechanisms to clinical trials. Front. Hum. Neurosci. 19, 1548478 (2025).

11. Bronte-Stewart, H. M. et al. Long-term personalized adaptive deep brain stimulation in Parkinson disease: A nonrandomized clinical trial: A nonrandomized clinical trial. JAMA Neurol. 82, 1171–1180 (2025).

12. Pedder, J. H. et al. Crossing the blood-brain barrier: emerging therapeutic strategies for neurological disease. Lancet Neurol. 24, 246–260 (2025).

13. Kadry, H., Noorani, B. & Cucullo, L. A blood-brain barrier overview on structure, function, impairment, and biomarkers of integrity. Fluids Barriers CNS 17, 69 (2020).

14. Ning, L., Makris, N., Camprodon, J. A. & Rathi, Y. Limits and reproducibility of resting-state functional MRI definition of DLPFC targets for neuromodulation. Brain Stimul. 12, 129–138 (2019).

15. Sun, W. et al. Precision network modeling of transcranial magnetic stimulation across individuals suggests therapeutic targets and potential for improvement. Hum. Brain Mapp. 46, e70266 (2025).

16. Fox, M. D. et al. Resting-state networks link invasive and noninvasive brain stimulation across diverse psychiatric and neurological diseases. Proc. Natl. Acad. Sci. U. S. A. 111, E4367–75 (2014).

17. Horn, A. et al. Connectivity Predicts deep brain stimulation outcome in Parkinson disease. Ann. Neurol. 82, 67–78 (2017).

18. Oathes, D. J. et al. Resting fMRI-guided TMS results in subcortical and brain network modulation indexed by interleaved TMS/fMRI. Exp. Brain Res. 239, 1165–1178 (2021).

19. Cash, R. F. H., Cocchi, L., Lv, J., Fitzgerald, P. B. & Zalesky, A. Functional magnetic resonance imaging-guided personalization of transcranial magnetic stimulation treatment for depression. JAMA Psychiatry 78, 337–339 (2021).

20. Weigand, A. et al. Prospective validation that subgenual connectivity predicts antidepressant efficacy of transcranial magnetic stimulation sites. Biol. Psychiatry 84, 28– 37 (2018).

21. Siddiqi, S. H. et al. Distinct symptom-specific treatment targets for circuit-based neuromodulation. Am. J. Psychiatry 177, 435–446 (2020).

22. Li, N. et al. A unified functional network target for deep brain stimulation in obsessive-compulsive disorder. Biol. Psychiatry 90, 701–713 (2021).

23. Horn, A. & Fox, M. D. Opportunities of connectomic neuromodulation. Neuroimage 221, 117180 (2020).

24. Laumann, T. O. et al. Functional System and Areal Organization of a Highly Sampled Individual Human Brain. Neuron 87, 657–670 (2015).

25. Gordon, E. M. et al. Precision Functional Mapping of Individual Human Brains. Neuron 95, 791–807.e7 (2017).

26. Braga, R. M. & Buckner, R. L. Parallel interdigitated distributed networks within the individual estimated by intrinsic functional connectivity. Neuron 95, 457–471.e5 (2017).

27. Seitzman, B. A. et al. Trait-like variants in human functional brain networks. Proc. Natl. Acad. Sci. U. S. A. 116, 22851–22861 (2019).

28. Gordon, E. M. et al. A somato-cognitive action network alternates with effector regions in motor cortex. Nature (2023) doi:10.1038/s41586-023-05964-2.

29. Dosenbach, N. U. F., Raichle, M. E. & Gordon, E. M. The brain’s action-mode network. Nat. Rev. Neurosci. 26, 158–168 (2025).

30. Hermosillo, R. J. M. et al. A precision functional atlas of personalized network topography and probabilities. Nat. Neurosci. 27, 1000–1013 (2024).

31. Thomas Yeo, B. T., et al. The organization of the human cerebral cortex estimated by intrinsic functional connectivity. J. Neurophysiol. 106, 1125–1165 (2011).

32. Gratton, C. et al. Functional Brain Networks Are Dominated by Stable Group and Individual Factors, Not Cognitive or Daily Variation. Neuron 98, 439–452.e5 (2018).

33. Demeter, D. V. & Greene, D. J. The promise of precision functional mapping for neuroimaging in psychiatry. Neuropsychopharmacology 50, 16–28 (2024).

34. Cole, E. J. et al. Stanford neuromodulation therapy (SNT): A double-blind randomized controlled trial. Am. J. Psychiatry 179, 132–141 (2022).

35. Lynch, C. J. et al. Automated optimization of TMS coil placement for personalized functional network engagement. Neuron 110, 3263–3277.e4 (2022).

36. Darrow, D. P., et al. An action networks model for pain reveals cortical neuromodulation targets. PsyArXiv (2025) doi:10.31234/osf.io/u4jky_v1.

37. Nahas, Z. et al. Prefrontal cortical stimulation for treatment-resistant depression. Brain Stimul. 18, 1334 (2025).

38. Hermosillo, R. et al. Convergence of individualized functional neural networks and structural connectivity in subdural prefrontal cortical stimulation. Brain Stimul. 18, 422 (2025).

39. Power, J. D. Resting-State fMRI: Preclinical Foundations. in fMRI 47–63 (Springer International Publishing, Cham, 2020).

40. Laumann, T. O. et al. On the stability of BOLD fMRI correlations. Cereb. Cortex 27, 4719– 4732 (2017).

41. Power, J. D., Barnes, K. A., Snyder, A. Z., Schlaggar, B. L. & Petersen, S. E. Spurious but systematic correlations in functional connectivity MRI networks arise from subject motion. Neuroimage 59, 2142–2154 (2012).

42. Power, J. D. et al. Methods to detect, characterize, and remove motion artifact in resting state fMRI. Neuroimage 84, 320–341 (2014).

43. Gordon, E. M., Laumann, T. O., Adeyemo, B. & Petersen, S. E. Individual variability of the system-level organization of the human brain. Cereb. Cortex 27, 386–399 (2017).

44. Fair, D. A. et al. Functional brain networks develop from a ‘local to distributed’ organization. PLoS Comput. Biol. 5, e1000381 (2009).

45. Dworetsky, A. et al. Probabilistic mapping of human functional brain networks identifies regions of high group consensus. Neuroimage 237, 118164 (2021).

46. Najafi, M., McMenamin, B. W., Simon, J. Z. & Pessoa, L. Overlapping communities reveal rich structure in large-scale brain networks during rest and task conditions. Neuroimage 135, 92–106 (2016).

47. Moore, L. A. et al. Towards personalized precision functional mapping in infancy. Imaging Neurosci (Camb) 2, 1–20 (2024).

48. Duprat, R. J. et al. Resting fMRI-guided TMS evokes subgenual anterior cingulate response in depression. Neuroimage 305, 120963 (2025).

49. Terao, I. & Kodama, W. Effectiveness of personalized repetitive transcranial magnetic stimulation for major depressive disorder: A systematic review and meta-analysis of randomized active-controlled trials. J. Affect. Disord. 381, 275–280 (2025).

50. Zarzycki, M. Z. & Domitrz, I. Stimulation-induced side effects after deep brain stimulation - a systematic review. Acta Neuropsychiatr. 32, 57–64 (2020).

51. Strotzer, Q. D. et al. Structural connectivity patterns of side effects induced by subthalamic deep brain stimulation for Parkinson’s disease. Brain Connect. 12, 374–384 (2022).

52. Lynch, C. J. et al. Frontostriatal salience network expansion in individuals in depression. Nature 633, 624–633 (2024).

53. Siddiqi, S. H. et al. Brain stimulation and brain lesions converge on common causal circuits in neuropsychiatric disease. *Nat*. Hum. Behav. 5, 1707–1716 (2021).

54. Ren, J. et al. Parkinson’s disease as a somato-cognitive action network disorder. Nature 651, 1030–1038 (2026).

55. Kaiser, R. H., Andrews-Hanna, J. R., Wager, T. D. & Pizzagalli, D. A. Large-scale network dysfunction in major depressive disorder: A meta-analysis of resting-state functional connectivity: A meta-analysis of resting-state functional connectivity. JAMA Psychiatry 72, 603–611 (2015).

56. Castellanos, F. X. & Proal, E. Large-scale brain systems in ADHD: beyond the prefrontal– striatal model. Trends Cogn. Sci. 16, 17–26 (2012).

57. Schilbach, L. et al. Transdiagnostic commonalities and differences in resting state functional connectivity of the default mode network in schizophrenia and major depression. NeuroImage Clin. 10, 326–335 (2016).

58. Doucet, G. E. et al. Transdiagnostic and disease-specific abnormalities in the default-mode network hubs in psychiatric disorders: A meta-analysis of resting-state functional imaging studies. Eur. Psychiatry 63, e57 (2020).

59. Zhong, X., Pu, W. & Yao, S. Functional alterations of fronto-limbic circuit and default mode network systems in first-episode, drug-naïve patients with major depressive disorder: A meta-analysis of resting-state fMRI data. J. Affect. Disord. 206, 280–286 (2016).

60. Shi, Y. et al. Abnormal functional connectivity strength in first-episode, drug-naïve adult patients with major depressive disorder. Prog. Neuropsychopharmacol. Biol. Psychiatry 97, 109759 (2020).

61. Hu, M.-L. et al. A review of the functional and anatomical default mode network in schizophrenia. Neurosci. Bull. 33, 73–84 (2017).

62. Liu, X., Zhang, N., Chang, C. & Duyn, J. H. Co-activation patterns in resting-state fMRI signals. Neuroimage 180, 485–494 (2018).

63. Gutierrez-Barragan, D., Ramirez, J. S. B., Panzeri, S., Xu, T. & Gozzi, A. Evolutionarily conserved fMRI network dynamics in the mouse, macaque, and human brain. Nat. Commun. 15, 8518 (2024).

64. Gutierrez-Barragan, D., Basson, M. A., Panzeri, S. & Gozzi, A. Infraslow State Fluctuations Govern Spontaneous fMRI Network Dynamics. Curr. Biol. 29, 2295–2306.e5 (2019).

65. Saggar, M., Shine, J. M., Liégeois, R., Dosenbach, N. U. F. & Fair, D. Precision dynamical mapping using topological data analysis reveals a hub-like transition state at rest. Nat. Commun. 13, 4791 (2022).

66. Shinn, M. et al. Functional brain networks reflect spatial and temporal autocorrelation. Nat. Neurosci. 26, 867–878 (2023).

67. Laumann, T. O., Snyder, A. Z. & Gratton, C. Challenges in the measurement and interpretation of dynamic functional connectivity. Imaging Neurosci. (Camb.) 2, imag–2–00366 (2024).

68. Boré, A., Guay, S., Bedetti, C., Meisler, S. & GuenTher, N. Dcm2Bids. (Zenodo, 2023). doi:10.5281/ZENODO.8436509.

69. Vizioli, L. et al. Lowering the thermal noise barrier in functional brain mapping with magnetic resonance imaging. Nat. Commun. 12, 5181 (2021).

70. Moeller, S. et al. NOise reduction with DIstribution Corrected (NORDIC) PCA in dMRI with complex-valued parameter-free locally low-rank processing. Neuroimage 226, 117539 (2021).

71. Dowdle, L. T. et al. Evaluating increases in sensitivity from NORDIC for diverse fMRI acquisition strategies. Neuroimage 270, 119949 (2023).

72. Esteban, O. et al. fMRIPrep: a robust preprocessing pipeline for functional MRI. Nat. Methods 16, 111–116 (2019).

73. Andersson, J. L. R., Skare, S. & Ashburner, J. How to correct susceptibility distortions in spin-echo echo-planar images: application to diffusion tensor imaging. Neuroimage 20, 870–888 (2003).

74. Glasser, M. F. et al. The minimal preprocessing pipelines for the Human Connectome Project. Neuroimage 80, 105–124 (2013).

75. Posse, S. et al. Enhancement of BOLD-contrast sensitivity by single-shot multi-echo functional MR imaging. Magn. Reson. Med. 42, 87–97 (1999).

76. Mehta, K. et al. XCP-D: A robust pipeline for the post-processing of fMRI data. Imaging Neuroscience 2, 1–26 (2024).

