## Supplemental Materials for "Precision Confidence Mapping: An approach to determining individualized network topography with limited data"

Funding:
J.S.B.R: T32 NIDA Training Grant [5T32DA007234-37];

B.T.C. K23DA057486

O.M.D. Research reported in this publication was supported by the University of Minnesota's MnDRIVE (Minnesota's Discovery, Research and Innovation Economy) initiative.

### Supplemental Materials

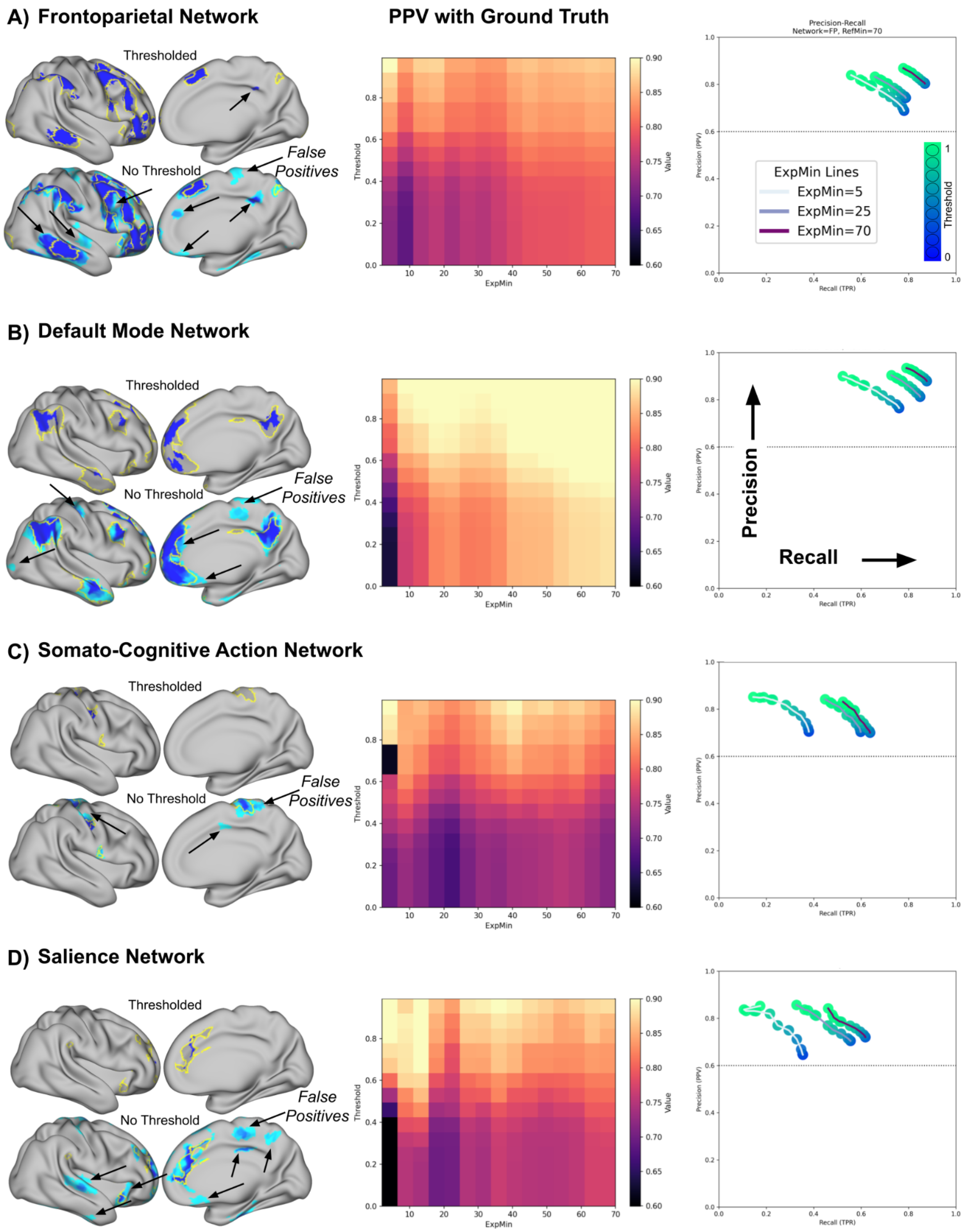

**Supplemental Figure 1.** Precision and TPR as a function of data length and confidence thresholding for networks commonly targeted in neuromodulation. **A)** Frontoparietal network. The top brain image shows a thresholded confidence map; the bottom shows a non-thresholded map. Arrows highlight regions of false positives that are present in the unthresholded map but are reduced after thresholding. The central heatmap depicts Positive Predictive Value (PPV), with the x-axis representing data length and the y-axis representing confidence map threshold. Thresholding improves PPV even at shorter data lengths. To the right, a precision–TPR plot demonstrates modest gains in targeting precision with increasing threshold, particularly at shorter scan durations. **B–D)** Same layout for the Default Mode Network (DMN), Somato-Cognitive Action Network (SCAN), and Salience network, respectively. The DMN exhibits the highest PPV and TPR across all networks, reflecting its robust and spatially consistent representation. In contrast, the SCAN and Salience networks display lower TPR (likely due to their smaller spatial extent), but maintain high precision, which further improves with confidence thresholding.

*Supplemental Table 1.*
*Effects of confidence thresholding on PPV and TPR across clinically relevant networks*

| Network | Duration | Metric | Thr = 0 | Thr = 0.99 | Δ | t(3) | p |
|---|---|---|---|---|---|---|---|
| DMN | 5 minute | PPV | 0.77 ± 0.04 | 0.91 ± 0.03 | 0.14 | 13.45 | < 0.001 |
| | | TPR | 0.76 ± 0.12 | 0.52 ± 0.18 | -0.24 | -6.79 | 0.007 |
| | 25 minute | PPV | 0.82 ± 0.08 | 0.91 ± 0.06 | 0.09 | 6.52 | 0.007 |
| | | TPR | 0.86 ± 0.10 | 0.74 ± 0.13 | -0.12 | -5.65 | 0.011 |
| | 70 minute | PPV | 0.88 ± 0.05 | 0.94 ± 0.04 | 0.06 | 10.45 | 0.002 |
| | | TPR | 0.88 ± 0.07 | 0.80 ± 0.09 | -0.09 | -5.56 | 0.011 |
| FPN | 5 minute | PPV | 0.69 ± 0.13 | 0.84 ± 0.11 | 0.16 | 9.72 | 0.002 |
| | | TPR | 0.79 ± 0.06 | 0.58 ± 0.10 | -0.22 | -10.70 | 0.002 |
| | 25 minute | PPV | 0.75 ± 0.20 | 0.84 ± 0.17 | 0.09 | 5.15 | 0.014 |
| | | TPR | 0.81 ± 0.12 | 0.69 ± 0.17 | -0.12 | -4.45 | 0.021 |
| | 70 minute | PPV | 0.80 ± 0.15 | 0.86 ± 0.12 | 0.06 | 4.78 | 0.017 |
| | | TPR | 0.88 ± 0.06 | 0.79 ± 0.08 | -0.08 | -5.43 | 0.012 |

| | | | | | | | |
|---|---|---|---|---|---|---|---|
| SCAN | 5 minute | PPV | 0.73 ± 0.10 | 0.85 ± 0.02 | 0.12 | 2.54 | 0.084 |
| | | TPR | 0.44 ± 0.21 | 0.18 ± 0.13 | -0.26 | -5.98 | 0.009 |
| | 25 minute | PPV | 0.71 ± 0.05 | 0.84 ± 0.03 | 0.13 | 4.79 | 0.017 |
| | | TPR | 0.66 ± 0.20 | 0.50 ± 0.18 | -0.16 | -5.50 | 0.012 |
| | 70 minute | PPV | 0.71 ± 0.10 | 0.84 ± 0.08 | 0.13 | 5.19 | 0.014 |
| | | TPR | 0.69 ± 0.19 | 0.57 ± 0.17 | -0.12 | -8.18 | 0.004 |
| SAL | 5 minute | PPV | 0.75 ± 0.21 | 0.88 ± 0.06 | 0.27 | 2.77 | 0.220 |
| | | TPR | 0.37 ± 0.17 | 0.18 ± 0.14 | -0.33 | -55.71 | 0.011 |
| | 25 minute | PPV | 0.75 ± 0.18 | 0.89 ± 0.11 | 0.14 | 3.74 | 0.033 |
| | | TPR | 0.59 ± 0.16 | 0.36 ± 0.16 | -0.23 | -11.81 | 0.001 |
| | 70 minute | PPV | 0.77 ± 0.20 | 0.88 ± 0.13 | 0.11 | 3.31 | 0.045 |
| | | TPR | 0.66 ± 0.19 | 0.50 ± 0.22 | -0.16 | -7.32 | 0.005 |
| All 15 networks | 5 minute | Mean PPV | 0.67 ± 0.04 | 0.86 ± 0.03 | 0.19 | 7.87 | 0.004 |
| | 25 minute | Mean PPV | 0.71 ± 0.04 | 0.85 ± 0.02 | 0.14 | 22.51 | < 0.001 |
| | 70 minute | Mean PPV | 0.75 ± 0.03 | 0.85 ± 0.02 | 0.10 | 14.15 | < 0.001 |

**Note.** Values are mean ± SD across subjects. Thr = 0 and Thr = 0.99 correspond to unthresholded and thresholded maps, respectively. Δ values were computed as threshold 0.99 minus threshold 0. Network abbreviations: DMN, default mode network; FPN, frontoparietal network; SCAN, somato-cognitive action network; SAL, salience network. The final rows report subject-level mean PPV averaged across all 15 canonical networks. Statistical comparisons were performed using paired two-sided t tests across subjects.

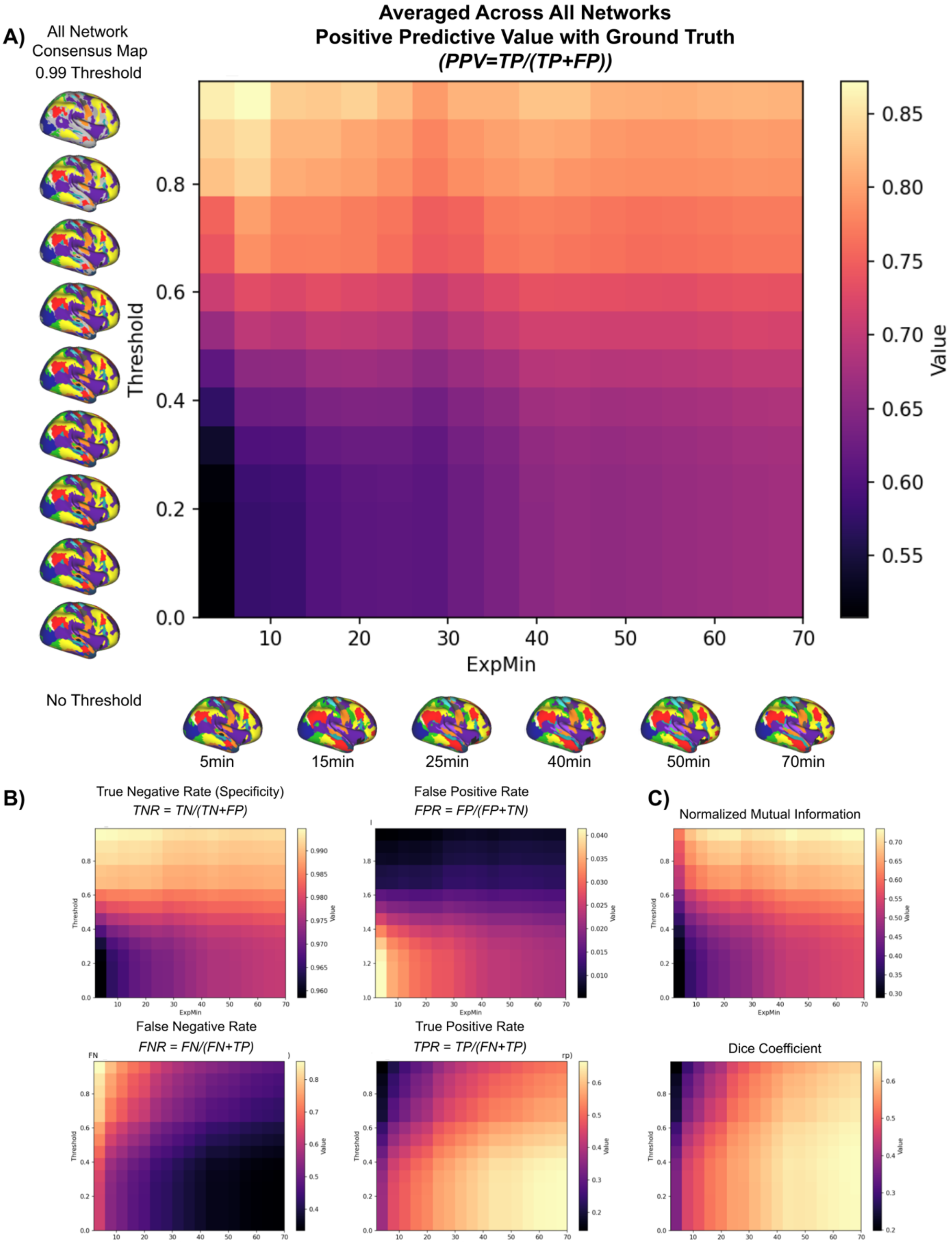
A)
All Network
Consensus Map
0.99 Threshold
Averaged Across All Networks
Positive Predictive Value with Ground Truth
(PPV=TP/(TP+FP))
Threshold
ExpMin
Value
No Threshold
5min
15min
25min
40min
50min
70min
B)
True Negative Rate (Specificity)
TNR = TN/(TN+FP)
False Positive Rate
FPR = FP/(FP+TN)
False Negative Rate
FNR = FN/(FN+TP)
True Positive Rate
TPR = TP/(FN+TP)
C)
Normalized Mutual Information
Dice Coefficient

**Supplemental Figure 2**. Confidence thresholding and increasing data length improve assignment accuracy across all canonical networks. A) Heatmap depicting Positive Predictive Value (PPV), calculated as TP divided by the sum of TP and FP, as a function of exploratory data length and confidence threshold. Values were calculated by comparing network assignments derived from exploratory Half 1 data with fixed reference assignments derived from 70 minutes of Half 2 data and were averaged across all 15 canonical networks. Representative whole brain network maps are shown alongside the axes to illustrate how data length and confidence thresholding affect the retained assignments. Heatmap color indicates mean PPV, which increases with confidence thresholding across data lengths, with the largest gains observed for shorter acquisitions. B) Complementary heatmaps show additional assignment metrics averaged across all 15 networks. The top row shows True Negative Rate on the left and False Positive Rate on the right. The bottom row shows False Negative Rate on the left and True Positive Rate on the right. Thresholding reduces false positive assignments and increases the True Negative Rate, with a corresponding increase in the False Negative Rate and modest reduction in the True Positive Rate. Longer data acquisitions generally improve performance across metrics. C) Normalized Mutual Information and Dice coefficient provide complementary measures of agreement between the exploratory and reference network assignments. The top heatmap shows NMI and the bottom heatmap shows Dice coefficient as functions of data length and confidence threshold. Together, these measures demonstrate how increasing data length and excluding lower confidence vertices affect agreement across whole brain network topography.

Supplemental Figure 3. PCM improves whole-network reliability across scan halves

The analyses in Figure 5 focused specifically on the action mode network (AMN) because a single network approach is most directly relevant to the neuromodulation applications motivating this work. To determine whether the reliability gains observed for the AMN generalized across the full set of canonical networks, we performed a complementary whole map analysis using normalized mutual information (NMI) to quantify within subject and between subject agreement across all assigned networks simultaneously. Unlike the fixed reference design used in Figure 5, this analysis compared independent halves of data of the same duration. It therefore provided a whole map assessment of test and retest reliability rather than an AMN specific assessment of targeting precision. We defined subject specificity as the extent to which within subject similarity exceeded between subject similarity, consistent with the comparison used in Figure 5D. For each participant, we calculated within subject NMI by comparing half 1 with half 2. We calculated between subject NMI as the mean similarity between that participant and all other participants. We then used paired tests across participants to determine whether within subject similarity exceeded between subject similarity. As in Figure 5, we compared standard template matching with PCM confidence maps thresholded at 0.99. Each comparison included only vertices assigned to a network in both halves.

With standard template matching and no PCM thresholding, within subject similarity exceeded between subject similarity but remained modest at 5 minutes. Within subject NMI was 0.42 ± 0.04, compared with a between subject NMI of 0.32 ± 0.01. The mean difference was 0.10 ±

0.03, with $t(3) = 5.90$ and $p = 0.0097$ across four participants. At 70 minutes, within subject similarity increased while remaining greater than between subject similarity. Within subject NMI was 0.66 ± 0.03, compared with a between subject NMI of 0.45 ± 0.01. The mean difference was 0.21 ± 0.02, with $t(3) = 26.48$ and $p = 1.18 \times 10^{-4}$. These findings were consistent with improved reliability at longer scan durations and preserved subject specificity, as shown in Supplemental Figure 3A.

PCM thresholding further increased within subject similarity while preserving the expected separation between within subject and between subject similarity. At 5 minutes with a confidence threshold of 0.99, within subject NMI increased to 0.84 ± 0.04, compared with a between subject NMI of 0.65 ± 0.01. The mean difference was 0.19 ± 0.05, with $t(3) = 7.81$ and $p = 0.0044$. At 70 minutes, PCM produced a within subject NMI of 0.88 ± 0.02, compared with a between subject NMI of 0.59 ± 0.02. The mean difference was 0.29 ± 0.01, with $t(3) = 74.66$ and $p = 5.30 \times 10^{-6}$, as shown in Supplemental Figure 3B.

Together, these results show that the improvement in within subject reliability produced by PCM while preserving subject specificity was not limited to the AMN. The same pattern was observed across whole brain network topography. This whole map analysis complements the AMN specific findings in Figure 5 and further supports the use of PCM for reliable individualized mapping in personalized neuromodulation applications.

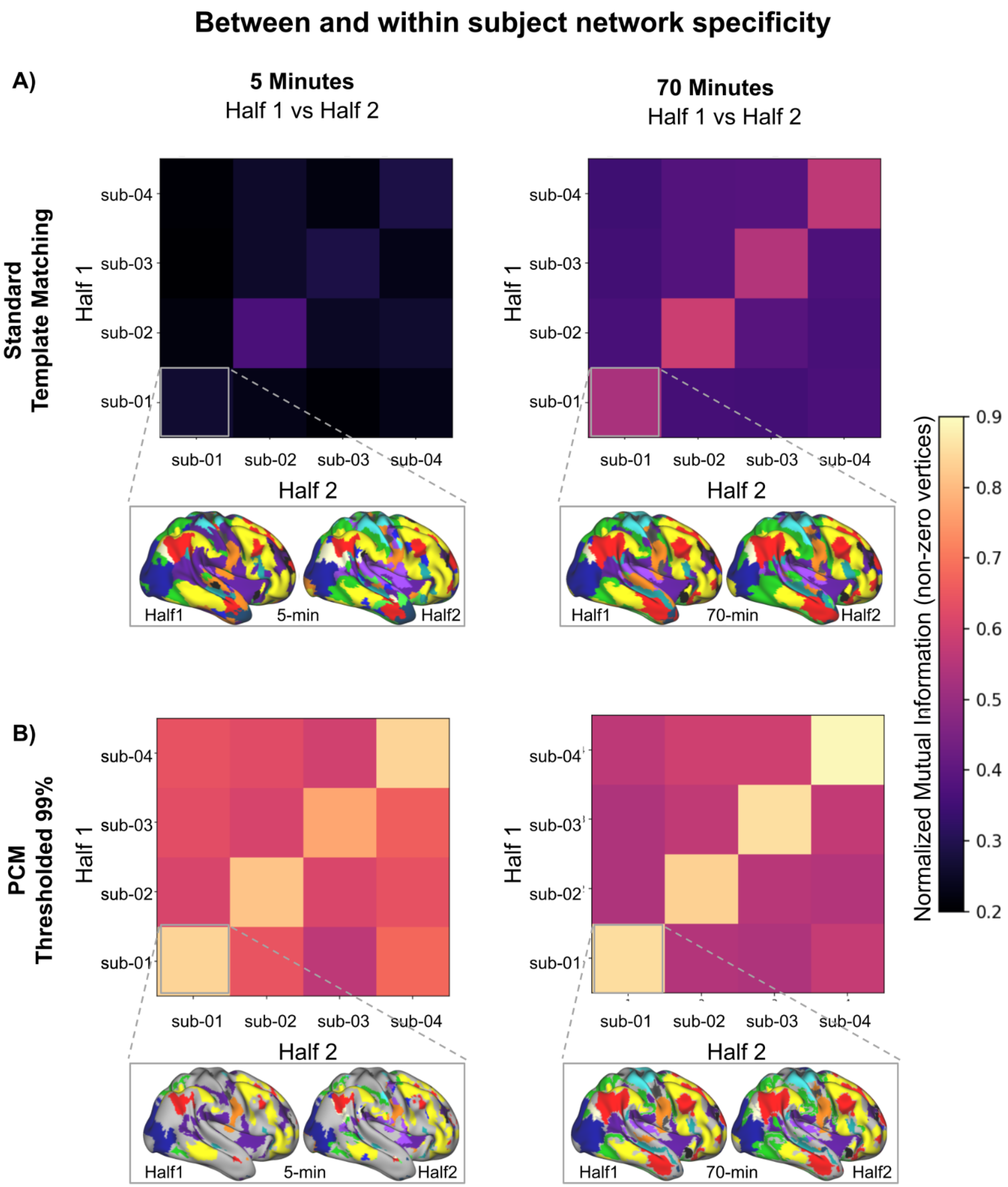

Between and within subject network specificity
A)
5 Minutes
Half 1 vs Half 2
70 Minutes
Half 1 vs Half 2
Standard
Template Matching
Half 1
sub-04
sub-03
sub-02
sub-01
sub-01
sub-02
sub-03
sub-04
Half 2
Half1
5-min
Half2
Half1
70-min
Half2
B)
PCM
Thresholded 99%
Normalized Mutual Information (non-zero vertices)
0.9
0.8
0.7
0.6
0.5
0.4
0.3
0.2

**Supplemental Figure 3**. PCM increases within-subject reliability while preserving subject specificity.

**(A)** Standard template matching network assignments were generated from 5 or 70 minutes of data for each subject using Half 1 and Half 2. Normalized Mutual Information (NMI) was computed using non-zero vertices only (vertices assigned to any network in both maps being compared) for all pairwise comparisons between Half 1 (rows) and Half 2 (columns). Diagonal entries show within-subject similarity (Half 1 vs Half 2 for the same subject); off-diagonal entries show between-subject similarity. At 5 minutes, within-subject similarity is modest and only slightly higher than between-subject similarity, whereas at 70 minutes within-subject similarity increases and remains higher than between-subject similarity, consistent with improved test–retest reliability while maintaining subject specificity. **(B)** The same analysis using PCM confidence maps thresholded at 0.99. Thresholded PCM maps show high within-subject NMI at both 5 and 70 minutes while preserving the within > between pattern, indicating that PCM enhances reproducibility without collapsing individual differences. Brain maps below each heatmap illustrate representative Half 1 and Half 2 network assignments at each scan duration (5 or 70 minutes).
